# Multiplexed quantitative screens of single cell shape and YAP/TAZ localisation identify DOCK5 as a coincident detector of polarity and adhesion during migration

**DOI:** 10.1101/2020.07.24.218313

**Authors:** Patricia Pascual-Vargas, Mar Arias-Garcia, Theodoros I. Roumeliotis, Jyoti S. Choudhary, Chris Bakal

## Abstract

YAP and TAZ are transcriptional co-activators that are often constitutively active in triple negative breast cancer (TNBC) cells driving proliferation, invasion, and drug resistance. Through multiplexed quantitative genetic screens for YAP/TAZ localisation and cell shape, we found that the RhoGEF DOCK5 is essential for YAP/TAZ activation in metastatic cells and is required for the maintenance of polarity during migration. DOCK5 regulates cell shape and thus YAP/TAZ through different genetic interactions with CDC42, RAC, and RHOA GTPases. DOCK5 regulates focal adhesion (FA) morphogenesis in RAC-dependent fashions that promote RHOA mediated actomyosin engagement of FA. Using unbiased systems-level quantification of protein levels by mass spectrometry we show that DOCK5 maintains polarity by stabilising protein levels of the CDC42 effector GSK3β. We conclude DOCK5 acts as a coincidence detector to promote leading edge persistence in subcellular locations where there is both RAC and RHOA dependent FA morphogenesis and active CDC42 mediated cell polarisation.

## Introduction

Many cells migrate by establishing a single, stable, polarised front, or leading edge (Etienne-Manneville, 2008; Ridley et al., 2003). The leading edge is the site of dynamic actin reorganisation that generates protrusive structures such as lamellipodia, and is the primary point of attachment to the extra-cellular matrix (ECM) by integrin-rich focal adhesions (FAs) (Etienne-Manneville, 2008; Ridley et al., 2003). Critically, despite constant turnover of both actin and integrin-based structures at the leading edge, and changes in ECM stiffness or geometry, many cells are able to maintain polarity for long periods of time and migrate in highly processive manners. However, how such polarity persists in migrating cells, despite large fluctuations in both their external environment and internal conditions, is unclear.

The activity of Rho-family GTPases is critically important to migration and to the formation of leading-edge structures. For example, the activation of CDC42 breaks symmetry and establishes polarity by activating atypical protein kinase C (PKCζ) which phosphorylates and inactivates GSK3β specifically at the leading edge. Phosphorylated GSK3β at the leading edge releases GSK3β from adenomatous polyposis coli protein (APC), which results in the association of APC at the cell cortex with microtubule (MT) plus-ends. Cortical APC binding captures MTs, which then results in reorganisation of the MT array establishing cell polarity(Etienne-Manneville, 2004; Etienne-Manneville and Hall, 2003a). Inactivated GSK3β determines sites of polarity in other cells such neurons (BARNES and POLLEUX, 2009; Gärtner et al., 2006; Shi et al., 2004; Zhou et al., 2004) and T-cells (Dong et al., 2013; Taylor and Rudd, 2020).

The ‘marking’ of cortical regions by activated CDC42 in migrating cells appears to be an evolutionarily conserved mechanism by which cells break symmetry (Johnson et al., 2011; Kozubowski et al., 2008). For example, activation of CDC42 in yeast cells initiates positive feedback loops by recruiting downstream Rho GTP Exchange Factors (Bos et al., 2007; Jaffe and Hall, 2005) such as CDC24 which then increase the levels of active CDC42 at the bud site (Freisinger et al., 2013; Goryachev and Leda, 2017; Wu et al., 2015). But how polarised structures initiated by CDC42 are maintained following their initiation in both yeast and mammalian cells remains unclear.

In migrating mammalian cells, the activation of RAC-type (RAC1, RAC2) GTPases at sites distal to the leading edge promotes FA growth, and reorganises actin (Hall, 1998; Ridley, 2001). Signalling by RHOA GTPase subsequently upregulates actomyosin tension which is key for the generation of traction at FAs (Case and Waterman, 2015; Gardel et al., 2010; Pertz et al., 2006; Ridley, 2012). But how the activity of Rho GTPases is regulated in migrating cells to coordinate FA dynamics and the maintenance of cell polarity is very poorly understood. The ability of some cancer cells to aggressively metastasise suggests that they have evolved mechanisms to dysregulate Rho GTPase activity in ways that favour processive migration and invasion (Clark et al., 2000; Fife et al., 2014; Jansen et al., 2018; Lawson and Ridley, 2018; Sahai and Marshall, 2002).

RhoGEFs have a number of protein-protein and protein-lipid interaction domains that mediate their recruitment to specific subcellular structures such as the membrane, nucleus, or FAs (Müller et al., 2020). At these specific milieus, RhoGEFs tightly regulate the spatiotemporal regulation of Rho GTPases, and even dictate the effectors that are recruited by Rho GTPases themselves (Jaffe and Hall, 2002, 2005). RhoGEFs can also act as non-enzymatic scaffolds that can act to assemble large signalling complexes (Rossman et al., 2005). Because of their ability to tightly regulate Rho GTPases in time and space, RhoGEFs are essential regulators of cell morphogenesis, migration and invasion in all eukaryotic cells (Bos et al., 2007; Fife et al., 2014; Fort and Blangy, 2017; Rossman et al., 2005).

RhoGEF mRNA and/or protein levels are frequently upregulated in breast cancer tissue (Adamowicz et al., 2006; Cook et al., 2014; Gatza et al., 2014; Lane et al., 2008; Laurin et al., 2013; Sahai and Marshall, 2002; Sosa et al., 2010). Further, the RhoGEFs *OBSCN* and *FGD5* have been found to be mutated in breast cancers (Cook et al., 2014; Gatza et al., 2014; Rossman et al., 2005; Shriver et al., 2015; Sjöblom et al., 2006). While the dysregulated activity of RhoGEFs has been implicated in driving the metastasis of cancer cells, and in particular breast cancer cells (Bravo-Cordero et al., 2011; Liu et al., 2009; Minn et al., 2005; Shriver et al., 2015), how RhoGEFs promote rapid changes in cytoskeleton, adhesion, and cell shape that drive processes essential to metastasis is largely unknown.

Increased RhoGEF and Rho GTPase activity has been linked to nuclear translocation of the transcriptional coactivators YAP and TAZ (hereafter referred to as YAP/TAZ). YAP/TAZ can promote a number of cancer relevant behaviours such as survival, proliferation, drug resistance, the Epithelial-to-Mesenchymal Transition (EMT), migration, and invasion (Cordenonsi et al., 2011; Dupont et al., 2011; Piccolo et al., 2014; Totaro et al., 2018). In normal cells, sequestration of YAP/TAZ proteins to the cytoplasm renders them inactive, and nuclear translocation of YAP/TAZ is trigged by soluble, mechanical, and geometrical cues (Aragona et al., 2013; Dupont et al., 2011; Lamar et al., 2012; Moroishi et al., 2015; Nallet-Staub et al., 2014; Pan, 2010; Sero and Bakal, 2017; Sero et al., 2015; Totaro et al., 2018; Zhao et al., 2008). Activation of YAP/TAZ is frequently observed in many cancer subtypes, and especially in breast cancer cells, although YAP/TAZ themselves are very rarely mutated in breast cancers (Cosmic; Harvey et al., 2013; Tate et al., 2019). Moreover, the evolved drug resistance of many cancers is due to dysregulation of YAP/TAZ activity (Calses et al., 2019; Fisher et al., 2017; Kapoor et al., 2014; Kim et al., 2016; Kitajima et al., 2018; Lin et al., 2015b; Nguyen and Yi, 2019; Shao et al., 2014). The mechanisms that lead to increased YAP/TAZ activity in cancers, and especially TNBCs are very poorly understood.

Activation of Rho GTPases promotes YAP/TAZ nuclear translocation in cancer cells through both direct and indirect means. Specifically, Rho GTPase activity alters the actin cytoskeleton resulting in YAP/TAZ activation in obscure fashions (Dupont et al., 2011; Piccolo et al., 2014; Totaro et al., 2018). Alternatively, Rho GTPases can recruit and activate enzymes such as PAK and ROCK kinases, which likely promote YAP/TAZ translocation directly (Sero and Bakal, 2017). Dysregulated RhoGEF activity can potentially explain YAP/TAZ activation in many cancer cells, and indeed oncogenic RhoGEFs such as TRIO have been found to promote YAP/TAZ nuclear translocation and activation (Feng et al., 2014, 2019). Finally, in cases where YAP/TAZ upregulation leads to evolved drug resistance, increased YAP/TAZ activation is almost always coincident with changes in cell shape, suggesting that changes in RhoGEF-Rho GTPase signaling could be drivers of this resistance by changing cell shape (Fisher et al., 2017; Kim et al., 2016; Lai et al., 2011; Lin et al., 2015a; Lionarons et al., 2019; Shao et al., 2014; Zanconato et al., 2016).

We sought to understand the basis for increased YAP/TAZ activity in metastatic breast cancer LM2 cells by performing multipliexed high-throughput genetic screens where we simultaneously measured YAP/TAZ nuclear translocation and cell shape following systematic knockdown of the majority of human RhoGEFs and RhoGAPs. These screens allowed us to identify genes important in both YAP/TAZ activation and cell shape, and led to the identification of the RhoGEF Dedicator of Cytokinesis 5 (DOCK5) as an essential regulator of YAP/TAZ activity in metastatic cells. Depletion of DOCK5 and lowered YAP/TAZ activity coincides with defects in migration following wound healing, impaired 3D invasion, and loss of drug resistance. To understand the basis for these phenotypes we explored how DOCK5 depletion affects the organisation of FAs and the cytoskeleton. We found that DOCK5 regulates the dynamics of FA morphogenesis, and the stabilisation of polarised leading edges, leading to dysregulation in cell shape determination. Using proteome-wide mass spectrometry approaches we show that the failure of DOCK5 depleted cells to maintain polarity is largely explained by significant decreases in GSK3β protein levels, and levels of GSK3β phosphorylation at the leading edge.

Mechanistically our results suggest that CDC42 activation seeds the formation of polarised structures independently of DOCK5, which involves phosphorylation of GSK3β. DOCK5 is recruited simultaneously to nascent FAs, where it acts in both RAC-dependent and independent fashions to promote FA morphogenesis and recruit more DOCK-AKT complexes in a positive feedback loop. DOCK5-AKT signalling stabilises GSK3β protein, which then sustains the establishment of polarity complexes by capturing FAs. Failure to stabilise GSK3β protein levels at the leading edge leads to an inability of microtubules to be captured at the leading edge – preventing the stabilisation of polarised structures.

Because DOCK5 acts at FAs to promote GSK3β stability at polarised fronts, this suggests that DOCK5 is a coincidence detector, or ‘AND gate’ which ensures polarity is maintained only where there is FA formation – i.e. at the leading edge of cells. By maintaining polarity only in FA rich regions, DOCK5 promotes adoption of polarised cell shapes that are capable of processive migration, sustained YAP/TAZ activation, and concomitant drug resistance. Our findings imply that cell behaviours underpinning oncogenesis and metastasis, such as drug resistance and invasion, can in part be driven by cells adopting polarised shapes, which in turn upregulates YAP/TAZ nuclear translocation.

## Results

### DOCK5 depletion lowers YAP/TAZ nuclear translocation

LM2 cells are a metastatic derivative of MDA-MB-231 TNBC cells (Minn et al., 2005). We have previously observed that in sub-confluent LM2 cells, YAP/TAZ proteins are predominantly nuclear (Pascual-Vargas et al., 2017). In fact, compared to the TNBC cell lines hs578t and SUM159, and the non-metastatic mammary carcinoma cell line T47D, LM2 cells had much higher nuclear YAP/TAZ at sub-confluent densities (Supplementary Fig. 1a). Increasing the cell density of LM2 cell populations to high levels (2.0 x 10^5^ cells per ml, 6000 cells per well) reduced YAP/TAZ nuclear translocation in LM2 cells, demonstrating that YAP/TAZ in LM2 cells is partially sensitive to cell numbers as observed in normal tissues (Sero and Bakal, 2017; Zhao et al., 2007) (Supplementary Fig. 1a). High basal levels of nuclear YAP/TAZ in sub-confluent LM2 cells also contrasted markedly to those observed in MCF10A normal breast epithelial cells (Supplementary Fig. 1a)(Sero and Bakal, 2017). Throughout this study we used LM2 cells at low density (3.3 x 10^4^ cells per ml, 1000 cells per well).

To identify regulators of Rho GTPases that contribute to the regulation of YAP/TAZ dynamics in TNBC cells, we systematically depleted 82 RhoGEFs, 67 GAPs and 19 Rho GTPases in LM2 cells and quantified the ratio of nuclear to cytoplasmic YAP/TAZ as a proxy for YAP/TAZ activation (Dupont et al., 2011; Sero and Bakal, 2017). LATS1, YAP and ECT2 siRNAs were used as positive controls (Pascual-Vargas et al., 2017): si*LATS1/2* for our ability to assess YAP/TAZ localisation, as YAP/TAZ nuclear localisation is expected to increase in the absence of LATS1/2 (Piccolo et al., 2013); si*YAP* for our ability to detect YAP/TAZ protein; and si*ECT2* to assess transfection efficiency as ECT2 depletion results in multinucleate cells (Miller and Bement, 2009; Su et al., 2011).

Knocking down individual Rho-family GTPases had distinct effects on YAP/TAZ nuclear translocation. Specifically, knockdown of CDC42, and RHOA resulted in a significant drop in nuclear YAP/TAZ (Fig. 1a). Knockdown of either RAC1 or RAC3 led to statistically insignificant decreases in nuclear YAP/TAZ, while knockdown of RAC2 led to a moderate decrease (Fig. 1a). Thus the basal levels of YAP/TAZ nuclear translocation observed in LM2 TNBC cells are partially due to the activation of CDC42 and RHOA. Though we cannot exclude the possibility that RAC1 and RAC2, but likely not RAC3 (Hajdo-Milasinović et al., 2007), promote YAP/TAZ nuclear translocation in LM2 cells in a partially redundant manner.

**Fig. 1.**
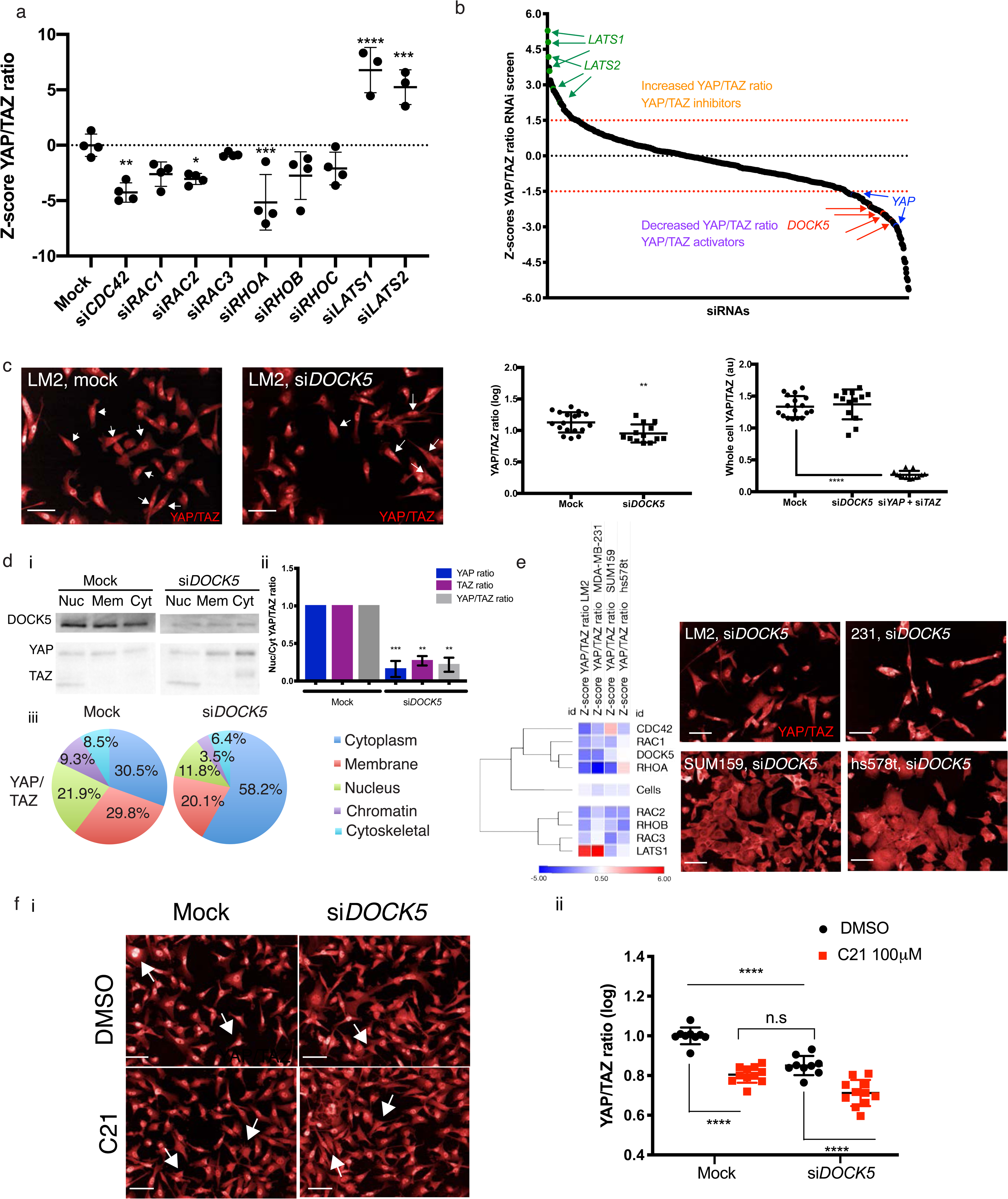
DOCK5 regulates YAP/TAZ localisation. (a) Graph depicting Z-scores of log YAP/TAZ ratios following knockdown of Rho GTPases. si*LATS1/2* included as controls for YAP/TAZ translocation. mean ± s.d (n = 3-4 wells per condition, 1000 cells per well, one-way ANOVA to mock). (b) Z-scores per well for YAP/TAZ ratio for OTP siRNA screen in LM2. Above 1.5 (red dotted line) are siRNAs which are hits for high YAP/TAZ ratio and thus can be considered inhibitors of YAP/TAZ. Below -1.5 (red dotted line) are siRNAs which are hits for low YAP/TAZ ratio and thus can be considered YAP/TAZ activators. *LATS1/2* and *YAP* siRNAs included as controls for high and low YAP/TAZ ratios are shown. Four individual replicates for si*DOCK5* are shown. (c) Representative images showing YAP/TAZ staining for mock transfected and si*DOCK5* transfected LM2 cells. Arrows point to cells with high YAP/TAZ ratio (mock) and low YAP/TAZ ratio (si*DOCK5*). Scale bar is 50μm. Graphs showing quantification of YAP/TAZ ratio, and whole cell YAP/TAZ levels in mock versus si*DOCK5.* si*YAP* + si*TAZ* show YAP/TAZ antibody is working appropriately. mean ± s.d (n = 3 independent biological repeats separate from screen data, dots represent individual wells, Student’s t-test (left) and ANOVA (right). (d) Validation of si*DOCK5* resulting in a decrease in YAP/TAZ ratio via subcellular fractionation. (i) Representative western blot of subcellular fractions for mock and si*DOCK5*. Nuc: nuclear fraction. Mem: membrane fraction. Cyt: cytoplasm fraction. Blotted for DOCK5, YAP and TAZ. Same amounts of protein loaded (6.5ng). (ii) Quantification of ratio of YAP/TAZ in the nucleus to the cytoplasm fraction from western blots of subcellular fractionations. mean ± s.d (n = 2). Student’s t-test. (iii) Representative distribution of YAP/TAZ in all fractions with and without *DOCK5*. (e) Heatmap for Z-score of YAP/TAZ ratio for knockdown of Rho GTPases and DOCK5 in OTP screens in TNBC cell lines LM2, MDA-MB-231, SUM159 and hs578t. Representative images for DOCK5 knockdown in each cell line. Scale bar is 50μm. (f) (i) Immunofluorescence images and (ii) quantification of YAP/TAZ localisation in mock or si*DOCK5* transfected cells treated with DMSO or DOCK5 inhibitor C21. mean ± s.d (n = 9 wells/condition, 1000 cells/well, one-way ANOVA). Significance for all panels: * P < 0.05, ** P < 0.001 *** P < 0.001, **** P < 0.0001. Scale bar is 50 μm.

To identify RhoGEFs that might be responsible for basal activation of YAP/TAZ in LM2 cells, we focused on RhoGEF siRNAs that reproducibly decreased nuclear YAP/TAZ levels. DOCK5 siRNA was the most consistent hit for siRNAs that caused low YAP/TAZ ratio (Fig. 1b), suggesting it is required for basal levels of YAP/TAZ activation in LM2 cells. 4 out of 4 replicates of DOCK5 siRNA were considered hits in our initial screen (Supplementary Table 1). We performed the same screen using another siRNA library (Dharmacon siGENOME; siG), where 4 out of 4 replicates of a DOCK5 siRNA pool with distinct sequences from the OTP pool also decreased YAP/TAZ nuclear levels (Supplementary Fig.1b, Z-scores per well: -1.73, -2.33, -2.77, -0.79). Notably, using both the OTP and siGENOME siRNA, depleting DOCK5 consistently reduced YAP/TAZ ratio without changing the total levels of YAP/TAZ (Fig. 1c, 1d, Supplementary Fig. 1c).

To validate that decreases in YAP/TAZ ratio were due specifically to DOCK5 depletion, we quantified mRNA expression levels following OTP and siGENOME knockdown of DOCK5 by quantitative RT-PCR and found that *DOCK5* mRNA expression levels were significantly reduced using either pool (Supplementary Fig. 1d). We also tested the ability of each individual sequence (4 siRNAs per pool) from the two different siRNA pools (OTP and siG) to knockdown *DOCK5* mRNA and found that all 8 siRNAs did so effectively (Supplementary Fig. 1e). Further, 7/8 single siRNAs significantly reduced YAP/TAZ ratio and none but siG3 reduced total YAP/TAZ levels (Supplementary Fig. 1c). Because it is virtually impossible that 7/8 DOCK5 siRNAs would reduce YAP/TAZ translocation through any other manner than by depleting DOCK5 mRNA and protein these data prove that DOCK5 promotes YAP/TAZ nuclear translocation in LM2 cells.

To corroborate our immunofluorescence imaging-based observations that DOCK5 depletion affects YAP/TAZ nuclear translocation, we performed subcellular fractionation of mock and DOCK5 depleted LM2 cells. In DOCK5 depleted cells the cytoplasmic fraction had significantly higher levels of both YAP and TAZ, compared to the same fraction from mock transfected cells. The ratios of nuclear to cytoplasmic YAP, TAZ, and the sum of YAP and TAZ (YAP/TAZ) were also significantly lower (Fig. 1d). The total amount of DOCK5 protein present in whole cell lysates for mock and si*DOCK5* transfected cells showed that DOCK5 protein was significantly reduced by DOCK5 siRNA (Supplementary Fig. 1g), indicating that DOCK5 siRNA is indeed specifically depleting DOCK5 as expected.

To understand whether the effect of DOCK5 depletion is specific to LM2 cells, or common across TNBC cell lines, we performed the same RNAi screens as for LM2, in the parental MDA-MB-231 cell line (∼ 1.5 million cells per screen, 4 wells per siRNA, 4000 cells/siRNA per screen), as well as two weakly metastatic TNBC cell lines: SUM159 and hs578t (∼ 2.5 million cells per screen, 4 wells per siRNA, 1500 to 2000 cells/well, 6000 - 8000 cells/siRNA per screen). We found that DOCK5 depletion also resulted in a YAP/TAZ translocation to the cytoplasm in both screens (siGENOME and OTP) in the parental cell MDA-MB-231 (Supplementary Fig. 2). Further, this decrease was also observed with RHOA siRNA in MDA-MB-231(Fig. 1e). However, DOCK5 was not a hit for YAP/TAZ translocation in SUM159 and hs578t cells (OTP, Fig. 1e). This suggests that DOCK5 plays a role in more strongly metastatic MDA-MB-231 and LM2 TNBC subtypes, but less so in weakly metastatic SUM159 and hs578t lines.

To test the role of DOCK5 RAC GEF activity on YAP/TAZ translocation we used C21, an inhibitor which blocks DOCK5’s DHR2 catalytic domain by hindering the interaction between DOCK5 and RAC1 (Ferrandez et al., 2017). Treatment with C21 decreased YAP/TAZ by the same amount as *DOCK5* siRNA (Fig. 1f). Thus DOCK5 catalytic activity on RAC-type GTPases is an essential component of how DOCK5 promotes YAP/TAZ nuclear translocation.

### DOCK5 contributes to growth and drug resistance in LM2 cells

In many cell types, YAP/TAZ activation is essential for survival and/or proliferation (Di Agostino et al., 2016; Lin et al.; Mizuno et al., 2012; Piccolo et al., 2013; Rosenbluh et al., 2012; Schlegelmilch et al., 2011; Shao et al., 2014; Yu et al., 2013). To determine if DOCK5, Rho GTPases, and basal YAP/TAZ activity contribute to the survival of LM2 TNBC cells we performed growth assays over 160 hours. We normalised confluency values per condition to that of mock transfected cells per well, and compared fold change to mock at 50% confluency (Methods). DOCK5 depleted cells grew at a slower rate as there were significantly less cells (68% less) following 60 hours post-transfection of DOCK5 siRNA where the mock population had reached 50% confluency (Fig 2a). Furthermore, DOCK5 depleted cells took 32 hours more than mock transfected cells to reach confluence (Fig. 2e, Supplementary Fig. 3a). Cells where CDC42 was depleted by siRNA also grew at a slower rate than mock transfected cells (Fig. 2a, b, Supplementary Figure 3b). But cells depleted of RHOA, RAC1, YAP, or TAZ grew at the same rate as mock transfected cells (Fig. 2a Supplementary Fig. 3c-f). However, the proliferation of cells simultaneously depleted of both YAP and TAZ was significantly impaired (Fig. 2a, Supplementary Fig. 3g). We conclude that DOCK5 and CDC42 promote proliferation of LM2 cells at least in part by promoting the nuclear translocation of both YAP and TAZ, though we cannot exclude the possibility that DOCK5 and/or CDC42 regulate proliferation independently of YAP/TAZ activation.

**Fig. 2.**
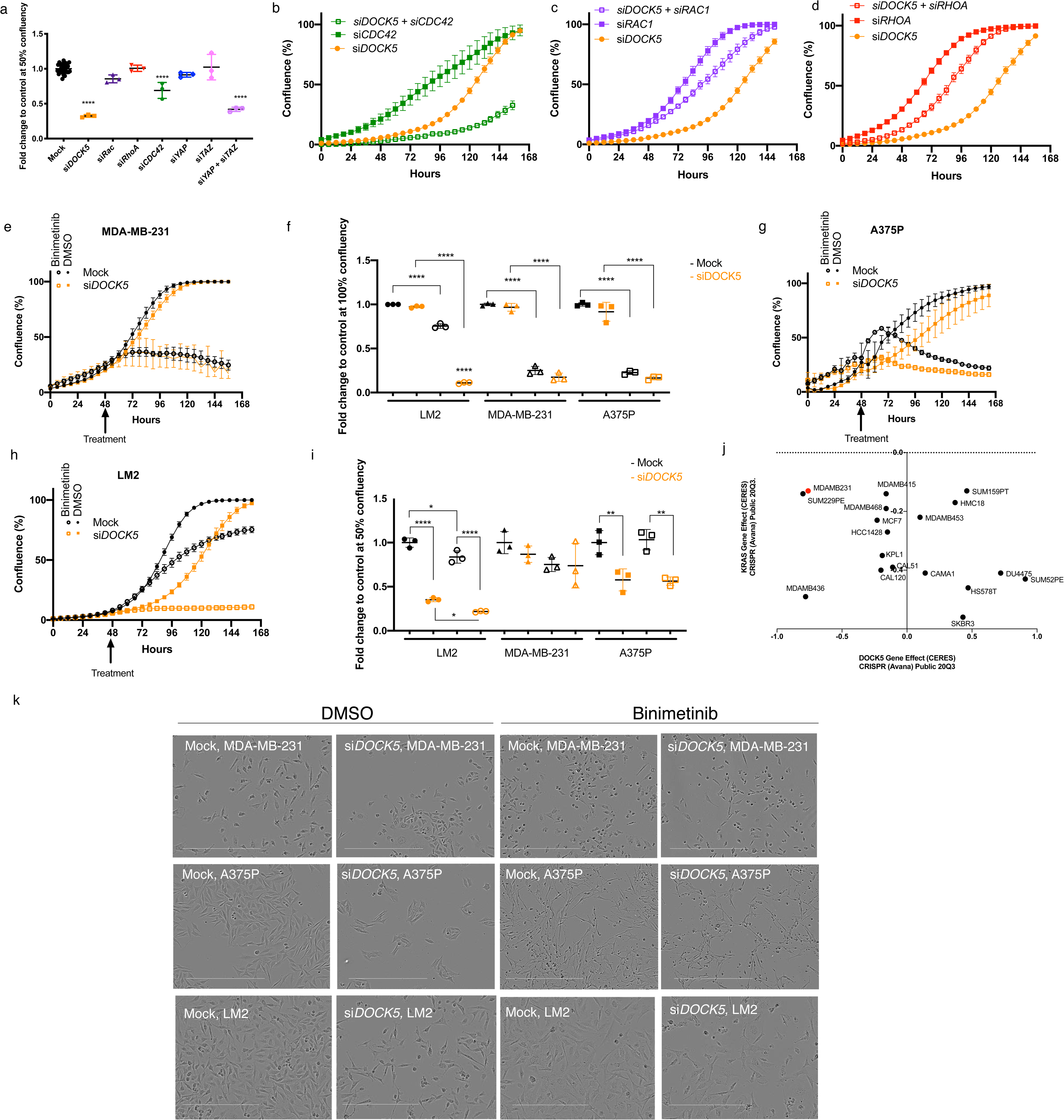
DOCK5 contributes to growth and drug resistance in LM2 cells. (a) Fold change to control mock transfected wells at 50% confluency for growth curves (Fig. 2b-d and Supplementary Fig. 3). Each data point is from one well. mean ± s.d, Student’s t-test. (b-d) Growth curves for (b) si*DOCK5*, si*CDC42*, si*DOCK5* + si*CDC42*, (c) si*DOCK5*, si*RAC1*, si*DOCK5* + si*RAC1,* (d) si*DOCK5*, si*RHOA*, si*DOCK5* + si*RHOA* transfected cells. Averages per well are shown. (e) (g) (h) Growth curves and (k) representative images at 50% confluency for mock and si*DOCK5* transfected cells treated with DMSO (closed circle, square) and 10 µM final MEKi Binimetinib (open circle, square) for (e) MDA-MB-231, (g) A375P, (h) LM2. (f, i) Fold change to control mock transfected wells at 100% and 50% confluency respectively for LM2, MDA-MB-231, and A375 mock and si*DOCK5* transfected cells treated with DMSO or Binimetinib. Each data point is from one well. mean ± s.d. Scale bar is 400 µm. All experiments performed in triplicate, representative experiment shown with n = 3 wells per condition. One-way ANOVA. For all panels: * P < 0.05, ** P < 0.01, *** P < 0.001, **** P < 0.0001. (j) Data plotted from Depmap website based on dependency of breast cancer cell lines to KRAS or DOCK5 knockdown. A CERES (computational method to estimate gene-dependency levels from CRISPR-Cas9 essentiality screens) (Meyers et al., 2017) a lower score indicates that cell line will be more dependent on that gene.

In order to determine if DOCK5 regulates survival in a fashion dependent on Rho GTPases, we assessed whether DOCK5 depletion is synthetically lethal with depletion of RAC1, RHOA, and CDC42. Indeed, DOCK5 and CDC42 exhibit synthetic lethality (Fig. 2b), such that at 156 hours when mock transfected cells were confluent, cells with both CDC42 and DOCK5 knocked down were at 32% confluency. In contrast, depletion of either RAC1 or RHOA with DOCK5 partially rescued the lethality of DOCK5 depletion (Fig. 2c, d respectively). We conclude that DOCK5 exhibits genetic interactions with CDC42, RAC1, and RHOA; a negative/aggravating interaction with CDC42 and positive/alleviating interactions with RAC1 and RHOA. Positive/alleviating interactions are symptomatic of “within pathway” interactions, which is consistent with the fact that DOCK5 activates RAC GTPase (Côté and Vuori, 2007; Laurin and Cote, 2014; Müller et al., 2020; Vives et al., 2011). That DOCK5 has a negative/aggravating interaction with CDC42 suggests they function independently of each other – i.e. in parallel pathways.

LM2 cells harbour mutant KRAS (G13D) (Lehmann et al., 2011), which activates the kinases MEK and ERK, driving proliferation and survival (Kumar et al., 2014; Tokumaru et al., 2020). However, while MEK inhibition (MEKi) effectively kills some cancer cells such as melanoma cells, breast cancer cells are often resistant to MEKi (Huang et al., 2017a; van der Noord et al., 2019). MEKi resistance in melanoma, pancreatic and non small lung cancer (NSLC) is often due to YAP/TAZ activation (Bado and Zhang, 2020; Coggins et al., 2019; Kitajima et al., 2018; Lin et al., 2015b). Therefore we determined whether the MEKi resistance of LM2 cells was due to DOCK5 and/or YAP/TAZ activity. To test this hypothesis we first treated LM2, MDA-MB-231 (from which LM2 cells are derived), and A375p melanoma cells (which harbour mutant BRAF) with the MEKi Binimetinib. MEKi significantly inhibited survival in MDA-MB-231 (Fig. 2e, f) and A375p (Fig. 2f, g) cells, but only had a modest effect on the survival of LM2 cells (Fig. 2f, h). DOCK5 depletion did not have any effect on the survival or proliferation of MDA-MB-231 or A375p cells in the absence or presence of MEKi (Fig. 2f, i). However, depletion of DOCK5 and MEKi treatment in LM2 cells led to a synergistic effect on proliferation or survival where confluency stalled at 10% for the whole duration at the assay (Fig. 2f, h, i). Thus, MEKi aggravates DOCK5 depletion suggesting that ERK and DOCK5 are functioning in compensatory pathways, and DOCK5 is essential for resistance of LM2 cells to MEK inhibition.

Mining of the DEPMAP database reveals that CRISPR-mediated depletion of DOCK5 and KRAS affects cell viability in a significantly similar manner across a large panel of cell lines (Fig. 2j; 0.4 Pearson correlation between effects of KRAS and DOCK5 depletion across cell lines as determined by DEPMAP profiling (Meyers et al., 2017)). Indeed, compared to all other breast cancer lines tested, MDA-MB-231 are most sensitive to either DOCK5 or KRAS CRISPR-mediated depletion. These data support the idea that both DOCK5-YAP and RAS-ERK pathways signalling promote survival in breast cancer cells, and that while inhibition of one axis can blunt survival, simultaneous inhibition of both is lethal.

### DOCK5 depletion reduces single cell invasion

To determine if DOCK5 contributes to the ability of LM2 cells to invade 3D tissues, we quantified the extent to which DOCK5 depleted cells invade rat tail collagen I (Col-I) gels. Cells typically invaded from the bottom of the plate into depths of 30μM in the collagen gel within 24 hours (Fig. 3a, b). We quantified single-cell invasion by calculating the invasion index, defined as the number of cells at various planes within the gel, divided by the total number of cells in all planes plus the bottom plane. In a 2.0 mg/ml rat-tail Col-I gel, DOCK5 depletion significantly reduced invasion (Fig. 3c). Further, depleting TAZ but not YAP significantly reduced invasion, by the same amount as depleting DOCK5. Thus TAZ drives LM2 invasion in 3D rather than YAP.

**Fig. 3.**
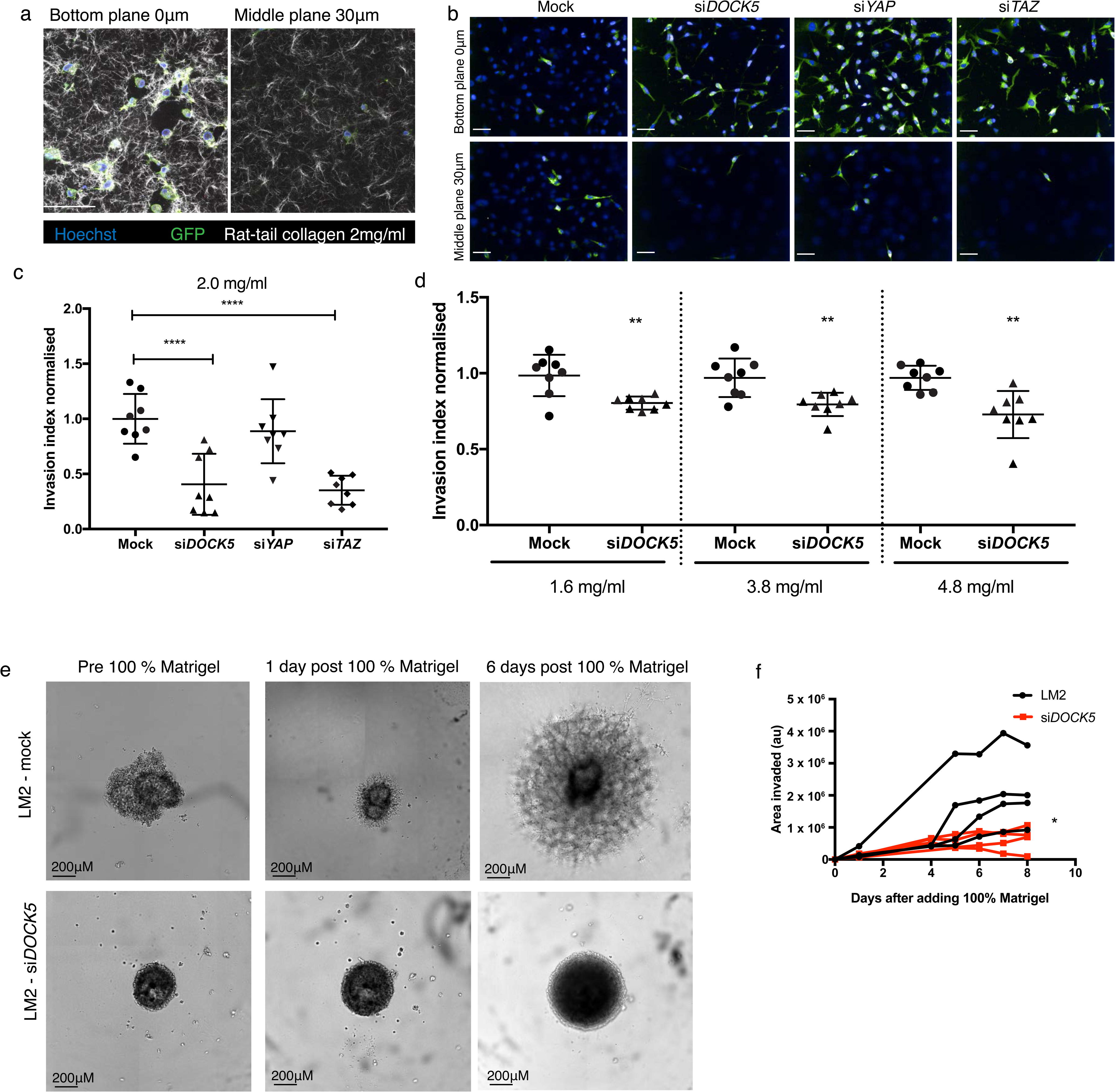
DOCK5 depletion reduces single cell invasion in 3D collagen assays and Matrigel spheroid assays. (a) Representative 2-photon images of LM2 cells (GFP) stained for DNA (blue) mixed with rat-tail collagen I at 2 mg/ml (white) at the bottom of the plate (0 µm plane) and the middle (30 µm plane). (b) Representative confocal spinning disk images (12 fields per well) for knockdowns shown in (c) at 0 µm and 30 µm. (c) Graph shows quantification of invasion index for knockdowns in rat-tail collagen I at 2 mg/ml. Invasion index is indicative of the number of cells in middle planes divided by total number of cells in all planes quantified, including bottom plane. Mean ± sd (n > 3 wells per condition, 1000 cells/well. ANOVA, **** P < 0.0001). (d) Invasion index of mock versus si*DOCK5* transfected cells at collagen concentrations: 1.6mg/ml, 3.8mg/ml, and 4.8mg/ml. Mean ± sd (n > 3 wells per condition, 1000 cells/well, ANOVA, ** P < 0.01). Scale bars, 100 µm. (e) Representative images of mock transfected cells and si*DOCK5* transfected cell spheroids 48 hours post transfection and pre 100% Matrigel (left), and one (middle) and six (right) days post 100% Matrigel addition. Scale bar is 200 μ m. (f) Graph shows quantification of area of Matrigel in arbitrary units over 8 days post adding 100% Matrigel to the spheroids. Mock transfected cells in black, si*DOCK5* transfected cells in red. Each line represents one well. n > 3 wells. Statistical significance obtained by performing student t-test on row means (* P = 0.0294).

Because collagen pore size has been shown to be inversely proportional to collagen concentration (Sapudom et al., 2015), we used collagen gels of different concentrations as a way of varying pore size. Depleting DOCK5 consistently reduced invasion across a Col-I density range of 1.6 to 4.8 mg/ml compared to mock transfected cells (Fig. 3d). Overall these data suggest that DOCK5 is required for LM2 single-cell invasion in rat tail collagen matrices independently of pore size.

To explore whether DOCK5 is required for invasion of cells in multicellular bodies – i.e. in tumour models, we generated spheroids and quantified cell invasion into 100% Matrigel (Methods). Mock transfected LM2 cells invaded into the gel progressively, while cells lacking DOCK5 did not invade at all (Fig. 3e,f). Overall this suggests that LM2 cells require DOCK5 to invade into 3D structures.

### DOCK5 depletion results in polarity and adhesion defects

We and others have shown that YAP/TAZ nuclear translocation is regulated by geometrical factors such as cell shape (Cordenonsi et al., 2011; Dupont et al., 2011; Pascual-Vargas et al., 2017; Sero and Bakal, 2017). Thus, we speculated that basal YAP/TAZ activation in LM2 cells and decreases in nuclear YAP/TAZ in DOCK5 depleted cells might be a consequence of cell shape changes. We sought to quantify how all RhoGEFs and RhoGAPs contribute to cell shape determination in metastatic TNBC cells. Using morphological profiling, we quantified the cell shape of single LM2 cells following depletion of 149 RhoGEF/GAPs and 19 Rho GTPases to generate quantitative morphological signatures or “QMSs” (Bakal et al., 2007). In this case, QMSs describe the percentage of cells with a particular shape in an RNAi-treated population, and thus account for the phenotypic heterogeneity observed in different cell lines (Yin et al., 2013). To generate QMSs, we measured 127 morphological and texture features for single cells in the screens, and trained linear classifiers to identify five visually distinctive reference shapes. The percentage of cells binned into each sub-population is a QMS for that specific RNAi (Methods). The five visually distinct shapes we used to characterize populations are: (1) ‘spindly’ - elongated cells with typically two protrusions; (2) ‘large round’-spread cells of a large area which are often circular; (3) ‘triangular’ - cells with three distinct protrusions; (4) ‘fan’ - asymmetrically-shaped cells with nucleus to one side; and (5) ‘small round’ - cells which have a small area and high roundness (Fig. 4a,b). LM2 cells are highly heterogeneous, and in a mock-transfected population 6.1% cells are classified as spindly, 1.1% as large round, 68.3% as triangular, 2.1% as fan, and 22.4% as small round (Fig. 4b, c). Depletion of different RhoGEF and RhoGAP genes by siRNA resulted in significant changes in the number of cells in each sub-population. For example, depletion of FAM13A resulted in 30.3% of cells being classified as spindly, 0.2% as large, 50.9% as triangular, 1.7% as fan and 16.9% as small round (Fig. 4d).

**Fig. 4.**
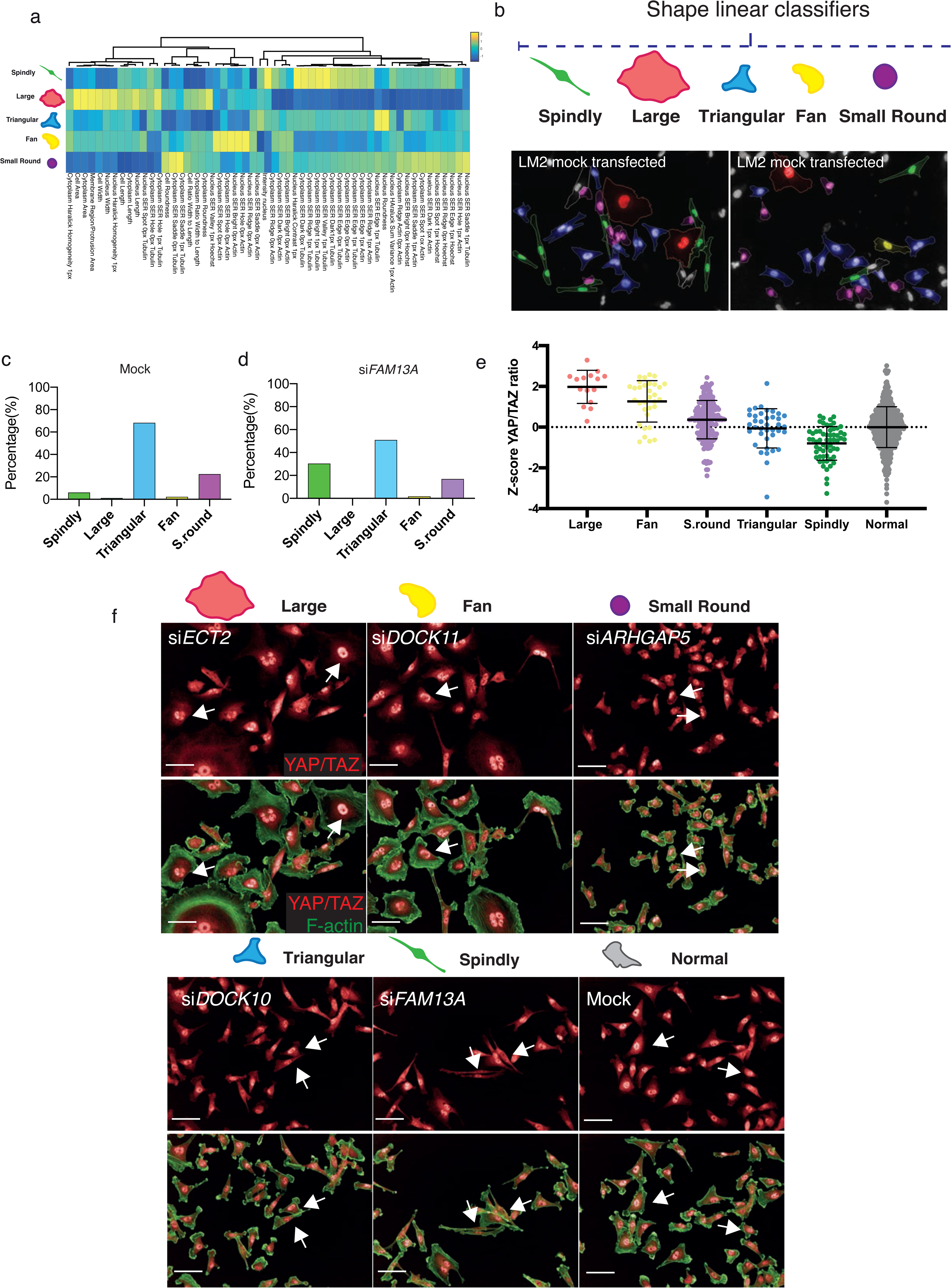
Shape classification and relationship with YAP/TAZ. (a) Heat map and dendrogram of the features describing each shape and which Columbus software used to classify cells into the five visually distinctive reference shapes, as well as their contribution, using single cell data. Yellow and blue represent a high and low score respectively in arbitrary units. (b) Images from one field of view from mock transfected well with cells coloured by their classified shape by software showing highly heterogenous population, and (c) the percentage of cells classified into each shape in one sample mock transfected well and (d) for an siRNAs which was a top hits for spindly shape, depicting a rearrangement in the shapes seen for mock-transfected cells. (e) Graph depicting Z-score of log10 ratio of nuclear to ring region YAP/TAZ for single LM2 cells following transfection with siRNA which results in the biggest enrichment for each particular shape: si*ECT2* for large, si*DOCK11* for fan, si*ARHGAP5* for small round, si*DOCK10* for triangular, si*FAM13A* for spindly. Mean ± sd are shown. Data taken from at least 5 fields of view from 2 wells. (f) Images of population from which single cell data shown in (e) were taken. Arrows indicate examples of nuclear YAP/TAZ for each shape. Images stained for YAP/TAZ (red) and F-actin (green). Scale bar is 50 μm.

We also developed a classifier for ‘normal’ cells (Methods, Supplementary Fig. 4). This allowed us to identify siRNAs which had significant effects on cell shape based on whether they significantly depleted the percentage of normal cells compared to mock (Z-score equal to or below -1 for normal classifier). In essence we could thus describe the penetrance of each genetic perturbation (Yin et al., 2013). 89 siRNAs resulted in significant depletion of normal cells; of which 79 were RhoGEF/GAPs, 7 were Rho GTPases, and 3 were controls for YAP/TAZ translocation si*LATS1*, si*LATS2* and si*YAP1*.

To determine if cell shape regulates YAP/TAZ in LM2 cells, we next asked whether exemplars of different cell shapes that were used to derive classifiers had predictable variability in YAP/TAZ nuclear translocation. Effectively asking the question, does shape predict YAP dynamics? Indeed, different classifier shapes had significantly different levels of nuclear YAP/TAZ ranging from high nuclear YAP/TAZ levels in polarised “fan” cells with broad lamellipodia to lower levels in “spindly” unpolarised cells (Fig. 4e, f). Thus as in other cell types (Dupont et al., 2011; Pascual-Vargas et al., 2017; Piccolo et al., 2014; Sero and Bakal, 2017; Totaro et al., 2018), in LM2 cells morphology can largely predict YAP/TAZ translocation dynamics.

To identify how different RhoGEFs, RhoGAPs, and Rho GTPases, and in particular DOCK5, regulate LM2 cell shape specifically, we performed hierarchical clustering on 5-shape QMSs for shape hits (<-1 Z-score for normal classifier) in the LM2 OTP screen, and defined “phenoclusters” as genes grouped at the highest node of clustering with a Pearson Correlation Coefficient (PCC) of > 0.73 (Fig. 5a, Supplementary Table 2). This resulted in 10 multigene clusters and 1 singleton formed by mock transfected LM2 cells.

**Fig. 5.**
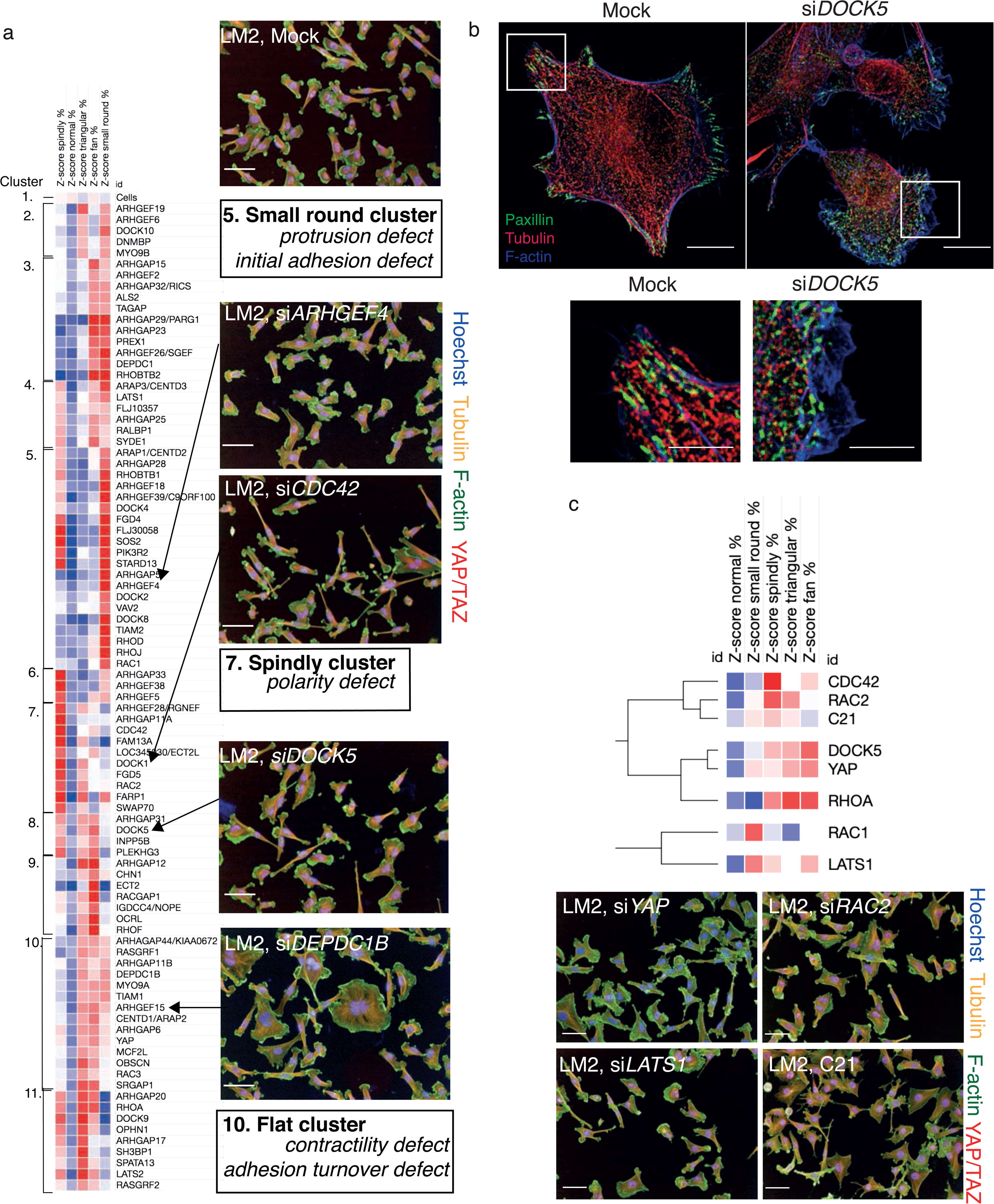
DOCK5 regulates shape. (a) (i) Clustering of high-dimensional quantitative signatures based on the effect of RhoGEF/GAPs and Rho GTPase siRNAs on the TNBC cell line LM2. Columns are Z-score values of the percentage of cells classified into that shape when said siRNA is depleted. Red and blue indicate high and low values respectively. Clustering performed using Pearson correlation. 10 clusters and a singleton were identified and defined as genes grouped at the highest node of clustering with a value of uncentered Pearson correlation coefficient above 0.73. Numbering indicates clusters and images are representative for mock and siRNAs which enrich for a particular shape in example clusters. Staining shows DNA (blue), tubulin (yellow), F-actin (green) and YAP/TAZ (red). Scale bar is 50 µm. (b) Representative images for a mock “normal” and an si*DOCK5* transfected cell imaged on SoRa. Staining shows Paxillin (green), α-tubulin (red), F-actin (blue). Square represents zoom in below. Scale bar is 10µm, zoom in 5µm. (c) Clustering of QMS for siRNAs shown as well as C21 treatment at a Pearson correlation of 0.74. Images for si*RAC2*, C21, si*LATS1*, and si*YAP* are shown, stained for DNA (blue), tubulin (yellow), F-actin (green) and YAP/TAZ (red). Scale bar is 50 µm.

The QMS of DOCK5 siRNA belonged to cluster 8 which is characterized by increases in fan, spindly, and triangular shapes (Fig 5a). DOCK5 depletion, and depletion of other genes in cluster 8, resembles spindle/unpolarised shapes following depletion of the canonical and conserved polarity regulator CDC42 (Cau, 2005; Etienne-Manneville and Hall, 2003a; Hall, 1994; Johnson and Pringle, 1990; Stowers et al., 1995) (cluster 7; Fig. 5a) and cluster 9 – which is characterised by very large flat cells such as following depletion of YAP/TAZ. We propose that the shape phenotypes of cluster 8 are likely due to defects in cytoskeletal organisation and/or FA assembly at the leading edge of polarized migrating cells. Indeed, DOCK5 depleted cells display disorganised microtubules which are not captured at the edge as well as lacking stabilised FA, contrary to mock “normal” cells where clear focal adhesions and microtubule capture can be observed (Fig. 5b). The similarity of CDC42, DOCK5, and YAP depletion on cell shape is consistent with our observations that both DOCK5 and CDC42 depletion individually decreased nuclear YAP/TAZ, and that DOCK5 and CDC42 exhibited synthetic lethality.

DOCK5 depletion affected cell shape determination in quantitatively similar ways as knockdown of: (1) CDGAP/ARHGAP31, a RhoGAP that localizes to FAs and regulates their dynamics during chemotaxis and durotaxis (He et al., 2011; LaLonde et al., 2006; Wormer et al., 2012), as well as regulating the actin cytoskeleton in integrin-independent ways (Jenna et al., 2002; Lamarche-Vane and Hall, 1998; McCormack et al., 2017); (2) PLEKHG3, an actin-binding protein that is essential for the establishment and maintenance of polarity in migrating cells (Nguyen et al., 2016); and (3) the endocytic regulator INPP5B (Festa et al., 2019; Williams et al., 2007) (Fig. 5a). Notably within cluster 8, only CDGAP/ARHGAP31 was also hit for low YAP/TAZ ratio in the OTP screen while PLEKHG3 and INPP5B had no effect (Supplementary Table 1). Thus while these data predict a role for all Cluster 8 genes in the regulation of polarity and focal adhesion dynamics, only CDGAP and DOCK5 can be considered YAP/TAZ regulators, and potentially part of a mechanism that promotes YAP/TAZ translocation by altering cytoskeleton organisation and cell shape.

We next asked whether the QMS of C21 treated cells is similar to that of DOCK5, YAP1, or LATS1 depleted cells. Because C21 inhibits GEF activity of DOCK5 towards RAC-type GTPases, we compared the DOCK5 QMS with QMSs following knockdown of CDC42, RAC1, RAC2, and RHOA. Hierarchical clustering of QMSs revealed that the QMS of C21 is most similar to RAC2 and CDC42 depleted cells (Pearson Correlation Coefficient (PCC) > 0.77), but only partially similar to the QMS of DOCK5 and YAP depleted cells (PCC > 0.40) (Fig. 5c). Thus DOCK5 depletion is not phenotypically identical to disruption of DOCK5 GEF activity, and suggests that DOCK5 mediated regulation of shape occurs both via GEF mediated regulation of RAC-type GTPase (i.e. RAC1 and/or RAC2), as well as by acting as a scaffold. That CDC42 knockdown mirrors the phenotype of C21 treatment, suggests that CDC42 activation may be partially reliant on DOCK5 catalytic activation of RAC-type GTPases, such that an effector of a RAC-type GTPases could potentiate the activity of active CDC42.

### DOCK5 depletion alters the levels of polarity proteins

In order to investigate how depletion of DOCK5 may affect FA dynamics, polarity establishment, and YAP/TAZ localisation, we used mass spectrometry to perform a proteome-wide analysis of total protein levels in mock and siDOCK5 transfected TNBC cell lines. Protein samples were processed 48 hrs after transfection and labelled with TMT-10plex (Tandem Mass Tag) multiplex reagents before being submitted to LC-MS/MS (liquid chromatography mass spectrometry; Methods). Protein abundance values obtained after analysis of mass spectra, scaled within cell lines, and the log2 ratio values of si*DOCK5* versus mock-transfected lines were calculated. DOCK5 protein was depleted by approximately 78% by siRNA (Fig. 6a, Supplementary Table 3). Further there were no changes in protein levels of DOCK5’s closest homolog, DOCK1 (Gadea and Blangy, 2014) (Fig. 6b), suggesting that the DOCK5 siRNA is not targeting DOCK1. In addition, there were no changes in protein levels of genes which clustered with DOCK5 by their QMS. Thus this confirms that DOCK5 siRNA has specific on-target effects and there is little chance of off-target effects.

**Fig. 6.**
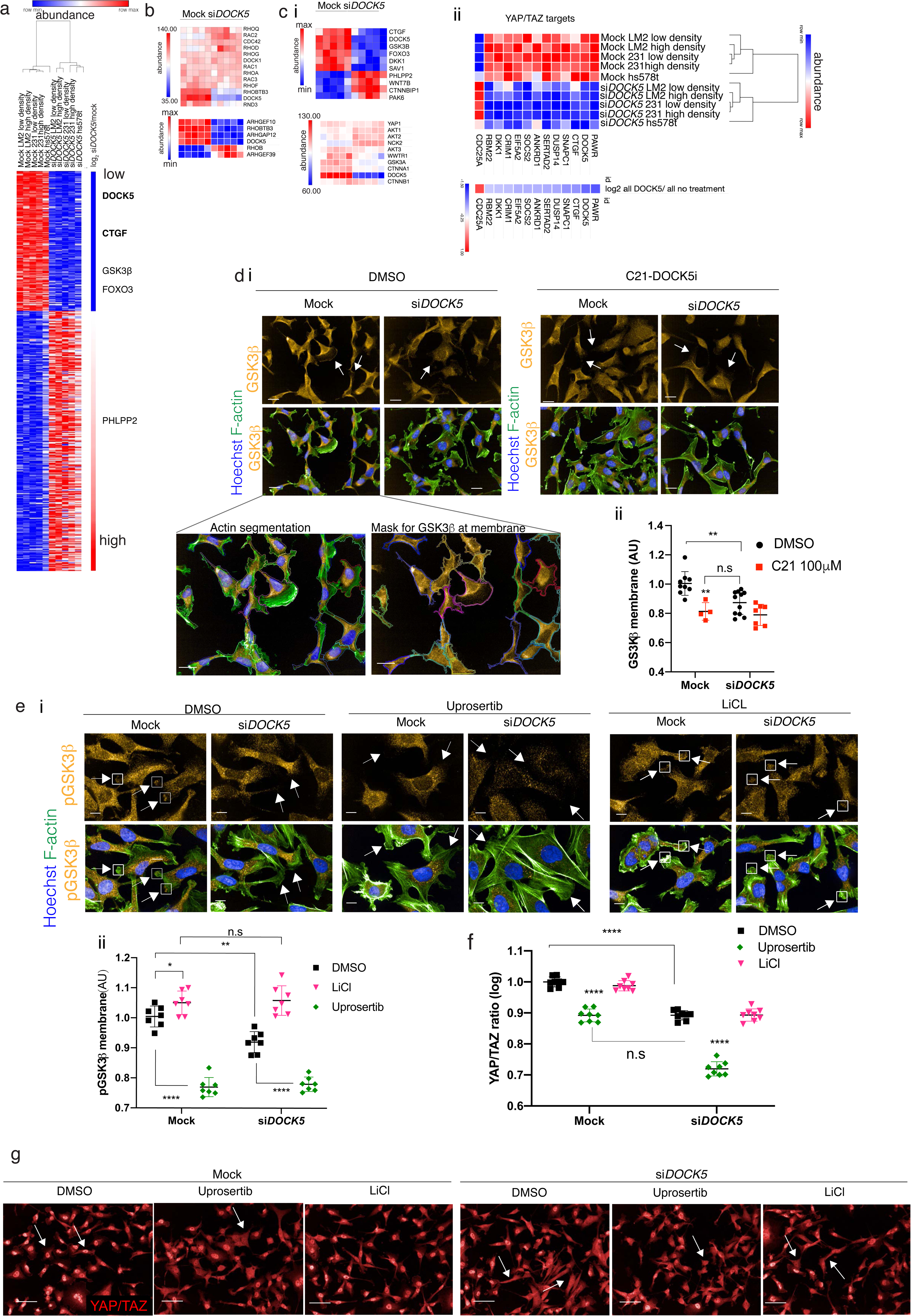
DOCK5 depletion alters protein levels of regulators of polarity. (a) Heat map of proteins significantly up or downregulated following *DOCK5* knockdown via siRNA in TNBC cell lines. Significance based on Log2 ratios for protein levels of si*DOCK5* v mock being above 0.5 and below -0.5 and permutation FDR <0.05 (t-test). Red and blue indicate high and low protein abundance levels (row max to min respectively). Proteins up and downregulated based on Log2 value. Proteins involved in regulating polarity and AKT signaling pathways shown are shown as well as CTGF. YAP/TAZ targets shown in bold (b) Top protein abundance levels of Rho GTPases and DOCK1 that were unchanged with si*DOCK5* abundance levels for comparison. Bottom protein levels of Rho GTPases and RhoGEF/GAPs which were significantly upregulated or downregulated upon DOCK5 knockdown. Row max (red) to min (blue). (c) (i) Top: Heat map of protein levels of proteins regulating polarity that were significantly up and down regulated. Values normalised to cell line. Row max (red) to min (blue). Bottom: abundance levels of proteins of interest that were not significantly up or downregulated, with si*DOCK5* abundance levels for comparison. (ii) Heat map of proteins significantly up or downregulated following *DOCK5* knockdown via siRNA in TNBC cell lines which have been shown to be YAP/TAZ target genes in MDA-MB-231. Red and blue indicate high and low protein abundance levels (row max to min respectively). Log2 value below. (d) (i) Representative images and (ii) immunofluorescence quantification of GSK3β at the membrane in mock and si*DOCK5* transfected cells in DMSO or DOCK5 inhibitor C21 Mean per well ± sd (n > 3 wells/condition, 1000 cells/well; ANOVA). White arrows indicate GSK3β at the membrane (mock) and lack thereof (si*DOCK5* and C21). GSK3β imaged at 40x, Scale bar, 20 µm. Inserts show actin segmentation and mask used to quantify GSK3β at the membrane. (e) (i) Representative images and (ii) immunofluorescence quantification of pGSK3β at the membrane in mock and si*DOCK5* transfected cells treated with DMSO (black), 5mM LiCl (pink) or 2μM final Uprosertib (green). Mean per well ± sd are shown (n > 3, 1000 cells/well, 2 ways ANOVA, Tukey’s multiple comparison’s test * P < 0.05, ** P < 0.01, *** P < 0.001, **** P < 0.0001). White arrows indicate pGSK3β at the membrane (mock and LiCl) and lack thereof (si*DOCK5* and Uprosertib). pGSK3β imaged at 60x with water objective (NA = 1.2). Scale bar, 10 µm. (f) Immunofluorescence quantifications of YAP/TAZ ratio of mock and si*DOCK5* transfected cells treated with DMSO (black), 5mM LiCl (pink) or 2μM final Uprosertib (green). Mean per well ± sd are shown (n > 3, 1000 cells/well, 2 ways ANOVA, Tukey’s multiple comparison’s test * P < 0.05, ** P < 0.01, *** P < 0.001, **** P < 0.0001). (g) Immunofluorescence images of YAP/TAZ for mock and si*DOCK5* transfected cells treated with DMSO (black), 5mM LiCl (pink) or 2μM final Uprosertib quantified in (f). Imaged at 20x. Scale bar, 50 µm.

Protein levels of Connective tissue growth factor (CTGF), which are upregulated by YAP/TAZ-TEAD activity (Zanconato et al., 2015; Zhao et al., 2008) were significantly downregulated (log2 ratio: -0.77695) (Fig. 6a, Supplementary Table 3); supporting the idea that depleting DOCK5 renders YAP/TAZ cytoplasmic and decreases YAP/TAZ transcriptional activity. Notably, total YAP and TAZ (encoded by the WWTR1 gene) protein levels remained unchanged (Fig. 6ci), matching our immunofluorescence data and confirming that DOCK5 regulates YAP/TAZ localisation rather than YAP/TAZ levels. In addition to CTGF, the protein levels of 10 established YAP/TAZ target genes (Zanconato et al., 2015) were also significantly downregulated following DOCK5 knockdown (Fig. 6c ii, Supplementary Table 3). These include SNAPC1 and PAWR, which have been shown to be positively regulated by YAP (Stein et al., 2015), further supporting the notion that DOCK5 alters YAP/TAZ localisation.

We performed hierarchical clustering on the abundance values of proteins which were significantly upregulated (40% increase) or downregulated (40% decrease) upon DOCK5 depletion (log2 ratios above 0.5 and below -0.5 and permutation FDR <0.05 (false discovery rate) (t-test)). DOCK5 depletion resulted in changes in the levels of AKT effectors, AKT regulators, and proteins implicated in the establishment of front-rear polarity. Specifically, the protein levels of the AKT effectors GSK3β (log2 ratio: -0.58753) and FOXO3A (log2 ratio: -0.59131) were downregulated (Fig. 6a, c). GSK3β plays a role in the establishment of polarity where it has been described to be phosphorylated and inhibited at the leading edge of migrating cells in a mechanism dependent on integrin mediated activation of CDC42-PAR6-PKCζ (Etienne-Manneville, 2004; Etienne-Manneville and Hall, 2003b, 2003a). Phosphorylated GSK3β promotes polarity by capturing/stabilizing microtubules (MTs) at the leading edge of migrating cells. Interestingly although it did not make the log2 ratio cut-off of -0.5 with a -0.46 log2 value, levels of the polarity regulator PAR6b were also substantially decreased (27%) in DOCK5 depleted cells (p = 0.0002, q = 0.0008). Finally, PHLPP2 which inhibits AKT (Brognard et al., 2007) and aPKCζ (Xiong et al., 2016), was upregulated (log2 ratio: 0.855787) (Fig. 6a, Supplementary Table 3). PHLPP2 upregulation has been shown to inhibit the establishment of polarity (Xiong et al., 2016). We conclude that depletion of DOCK5 results in decreased levels of signalling proteins and activity that maintain polarity, and is consistent with increased numbers of unpolarised, especially spindly-shaped cells following DOCK5 depletion (Fig. 5). Increasing spindly-shaped cells, which have low YAP/TAZ activation, likely explains the bulk of overall decreases in YAP/TAZ activity following DOCK5 depletion.

DOCK5 has been shown in osteoclasts to promote serine 9 (Ser9) phosphorylation downstream of AKT activation in both RAC dependent and independent ways that lead to MT stabilisation (Guimbal et al., 2019; Ogawa et al., 2014). To investigate whether DOCK5 regulates the levels of phosphorylated GSK3β at the leading edge to promote MT stability and the maintenance of cell polarity, we used quantitative single-cell imaging approaches to quantify the subcellular localisation of GSK3β at the tips of cell protrusions (Fig. 6d). Validating the observation that DOCK5 depleted cells have significantly decreased levels of GSK3β as determined by mass spectrometry, DOCK5 depleted cells had significantly reduced levels of GSK3β at the edge of protrusions compared to control cells (Fig. 6d). Total GSK3β levels were also reduced by C21 treatment, demonstrating that DOCK5’s exchange activity on RAC-type GTPases is essential to maintain GSK3β levels in polarised protrusions. We propose that the inability of DOCK5 depleted cells to recruit GSK3β to the membrane leads to defects in the maintenance of polarity.

To determine the role of DOCK5 and AKT in regulating levels of phosphorylated Ser9 GSK3β at protrusions we quantified the levels of phosphorylated GSK3β in LM2 cells at membrane regions following DOCK5 depletion and/or treatment with the AKT inhibitor Uprosertib (final concentration 2μM). We confirmed that Uprosertib indeed inhibits AKT by measuring FOXO3A nuclear translocation (Supplementary Fig. 5a, b) (Brunet et al., 2001; Manning and Toker, 2017). We also used LiCl (final concentration 5mM) as a positive control of our ability to measure GSK3β phosphorylation (pGSK3β). As expected, LiCl caused an increase in pGSK3β levels in protrusions in mock-transfected cells due to disruption of the negative feedback on AKT-mediated Ser9 phosphorylation by GSK3β kinase activity (O’Brien et al., 2011). Further, we confirmed LiCl led to a significant increase in whole cell β-catenin levels via immunofluorescence (Supplementary Fig. 5c, d) (Erdal et al., 2005; Huang et al., 2017b; Schleicher et al., 2017; Wu and Pan, 2010).

DOCK5 depleted cells significantly decreased pGSK3β (Etienne-Manneville and Hall, 2001) at the leading edge, while AKT inhibition resulted in an even greater decrease in pGSK3β levels at the protrusive edges in both mock and si*DOCK5* transfected cells (Fig. 6e). Strikingly, LiCl treatment rescued the phenotype of pGSK3β levels in DOCK5 depleted cells resulting in the same levels of pGSK3β in the protrusions of LiCl treated si*DOCK5* and mock transfected cells (Fig. 6e). The ability of LiCl to restore pGSK3β levels in DOCK5 depleted cells is likely because LiCl blocks GSK3β’s ability to inhibit AKT mediated phosphorylation of Ser 9 independently of DOCK5 (O’Brien et al., 2011).

Taken together these data support a mechanism by which DOCK5 acts as a RAC GEF to promote GSK3β stability and phosphorylation by AKT and other kinases. Combined with prior observations that DOCK5 acts to recruit AKT to different subcellular milieus (Guimbal et al., 2019; Ogawa et al., 2014), we propose that recruited AKT is likely activated downstream of RAC to promote GSK3β Ser9 phosphorylation and GSK3β stability.

In order to test whether YAP/TAZ translocation to the cytoplasm could be due to decreases in the levels of stable phosphorylated GSK3β at the membrane, and decreased AKT signalling, we quantified YAP/TAZ translocation following inhibition of AKT or GSK3β kinase activity, in both mock and si*DOCK5* treated cells. Inhibiting AKT directly with Uprosertib resulted in the same decrease in YAP/TAZ ratio in mock transfected cells similar to depleting DOCK5 by RNAi (Fig. 6f, g). Furthermore, treating DOCK5 depleted cells with Uprosertib decreased the ratio significantly compared to mock transfected cells (Fig. 6f, g). LiCl however had no effect on YAP/TAZ. Thus elevating pGSK3β levels in control cells at the membrane alone does not further increase YAP/TAZ translocation. Hence, we propose a DOCK5-RAC-AKT complex contributes to YAP/TAZ activation in LM2 cells. We cannot determine if the effects of DOCK5-RAC-AKT signalling directly tactivate YAP/TAZ - for example via chemical signalling pathways (Kim and Gumbiner, 2015; Sero and Bakal, 2017) - or indirectly by affecting focal adhesion morphogenesis and cell shape.

### DOCK5 depletion reduces wound closure rates and affects Golgi orientation

To test our hypothesis that DOCK5 contributes to the establishment of front-rear polarity in migrating cells, we performed wound healing assays, which are tests of polarity initiation and maintenance, 2D migration speed, and migration persistence (Methods). We also examined whether YAP/TAZ activation may be important for polarity establishment and migration. In these assays, we depleted DOCK5, YAP, TAZ, and YAP and TAZ in combination. Three time points from the 16 hr assays are shown in Fig. 7a for all conditions. Depleting DOCK5, TAZ, and YAP + TAZ significantly reduced migration rates but depleting YAP did not (Fig. 7a, b). Thus, DOCK5 and TAZ promote migration in 2D.

**Fig. 7.**
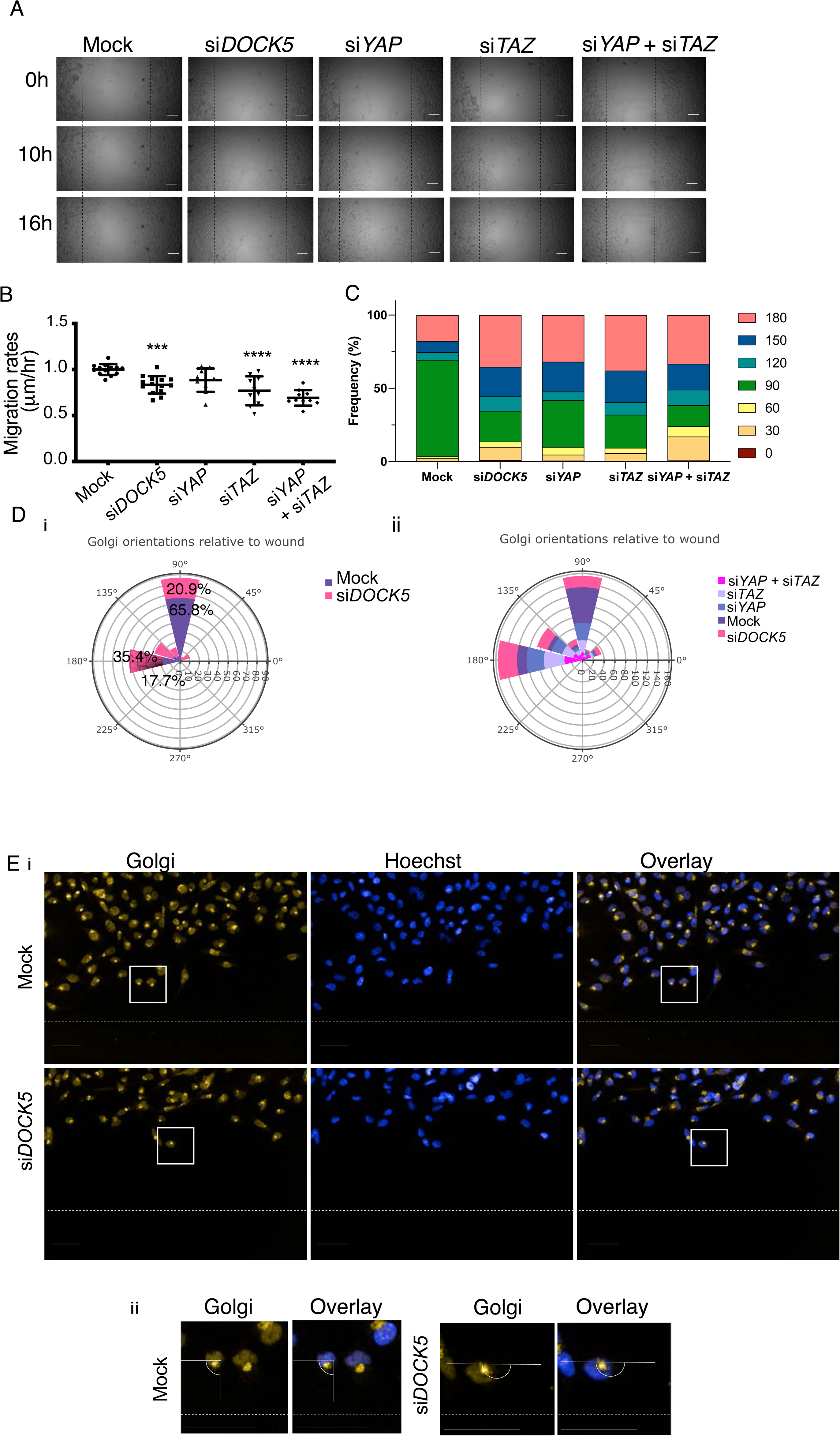
DOCK5 depletion reduces wound closure migration rates and affects Golgi orientation. (a) Brightfield images at 0, 10 and 16 hours post creating scratch wound for mock, si*DOCK5*, si*YAP*, si*TAZ*, and si*YAP* + si*TAZ* transfected cells. Dotted black line indicates the starting line from where cells migrated over time. Scale bar, 100 µm. (b) Graph shows migration rates (µm/hour) (n = 5 wells per condition, with at least 2 biological repeats). Values were normalised to average of mock for each biological repeat. Mean ± sd, one-way ANOVA to mock. *** P < 0.001, **** P < 0.0001 (c), (d) Percentage frequency of cells with Golgis orientated 0, 30, 60, 90, 120, 150, and 180 degrees relative to the wound depicted as a bar chart (c) and as a windrose plot (di,ii). Percentages indicated on windrose plots. (e) Images depicting Golgi immunofluorescence stain and Hoechst, wound is shown with white dotted line. Angle measuring relative to wound is depicted in (ii), example of 90 degree angle and orientation towards the wound. Scale bar is 50 µm.

The establishment and maintenance of polarity during wound healing is characterised by the reorientation of the Golgi, and microtubule organising centre (MTOC), toward the wound area and direction of migration (Etienne-Manneville and Hall, 2001). Therefore to determine if DOCK5 contributes to the initiation and/or establishment of leading edge polarity, we quantified the orientation of the Golgi following wounding, in cells at the edge of the wound (Etienne-Manneville, 2004; Zhang et al., 2008). Specifically, we quantified the angle of the Golgi oriented to the wound such that a 90 degree angle indicates the Golgi is facing toward the wound and front-rear polarity has been established (Fig. 7e, insert). Mock transfected populations had a high frequency of cells with Golgi directly facing the wound (65.8%) (Fig. 7c, di). In contrast, DOCK5 depleted populations had a higher frequency of Golgi oriented parallel to the wound (180 degrees) and/or that were poorly polarised (35.4% to 17.7% respectively) (Fig. 7c, di). YAP and TAZ combined depletion resulted in the condition with the least percentage of cells displaying front-rear polarity (with Golgi oriented 90 degrees to the wound) and the highest percentage of cells with Golgi oriented 30 degrees relative to the wound, suggesting these cells are unable to integrate cues from the microenvironment for directional motility (Fig 7c, dii). Taken together, these data support the idea that DOCK5 promotes the formation of polarised leading edge protrusions.

### DOCK5 depletion prevents the maintenance of polarity and alters FA dynamics

We and others have demonstrated that DOCK5 localises in close proximity to FAs peripherally to Paxillin, but co-localises with Paxillin in the absence of GIT2 (Frank et al., 2017; Müller et al., 2020). In line with these data, we observed that overexpressed DOCK5-YFP is also localised in puncta at the very edge of the lamellipodial protrusions in TNBC cells. DOCK5 bordered the forward-facing side of Paxillin-positive FAs at the leading edge (Fig. 8a). Notably, the rear facing side of Paxillin-positive FAs at the leading edge was often bordered by polymerised alpha-tubulin (Fig. 8a).

**Fig. 8.**
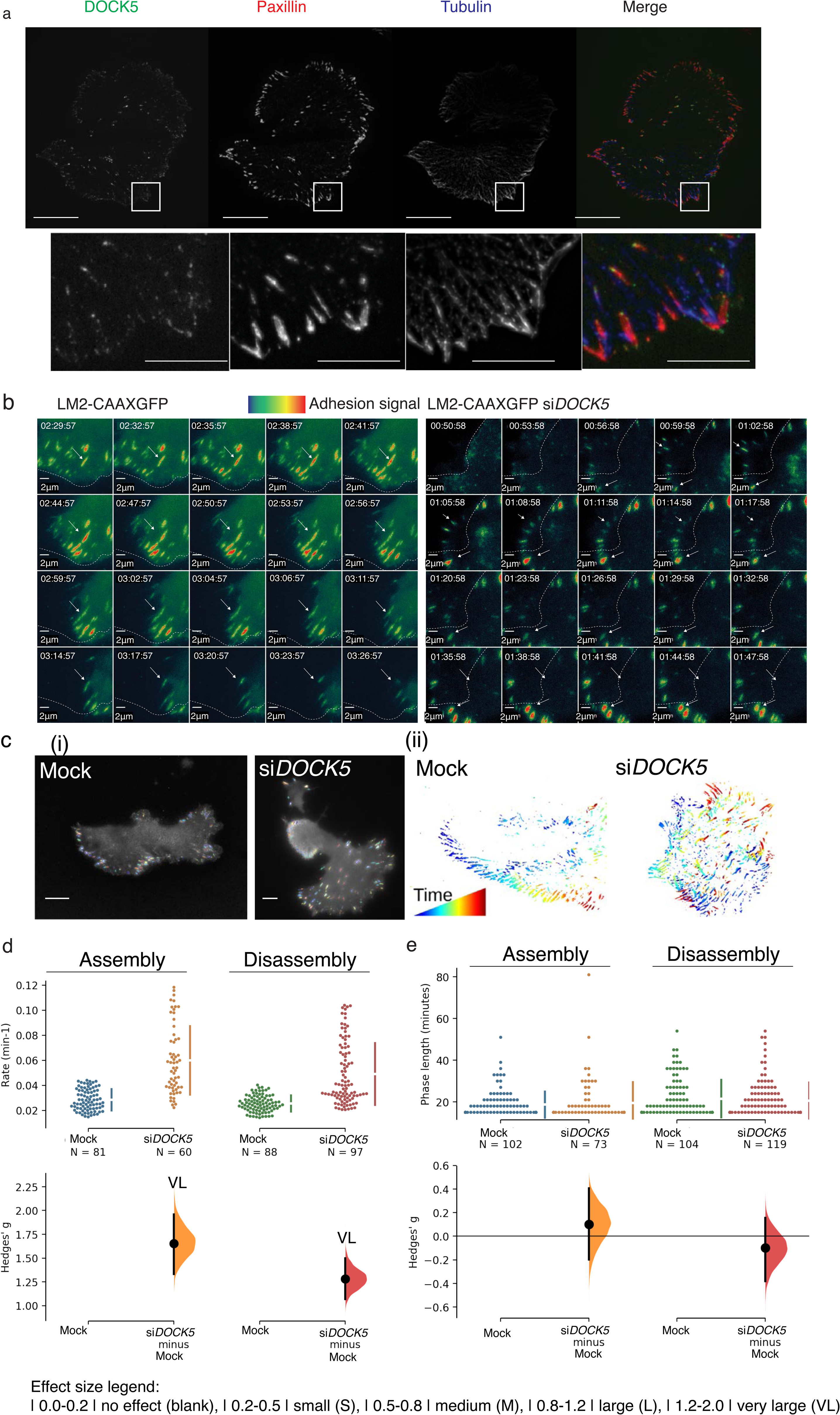
DOCK5 depletion prevents the maintenance of polarity and alters focal adhesion dynamics. (a) Overexpressed DOCK5-tagged YFP localises to focal adhesions (paxillin). Cells stained for paxillin (red), tubulin (blue). Scale bar is 5 µm. (b) Sequences of zoomed in sections of mock, and si*DOCK5* transfected LM2-CAAXGFP cells over time. Frames are every 3 minutes. Arrows indicate an example of an adhesion to follow. Dotted line represents the edge of the cell at the beginning of each sequence of frames shown. Adhesion signal represents Talin mApple intensity: red more intense, blue less intense. Scale bar is 2 µm. (c) (i) Representative images of how Focal Adhesion Analysis Server (FAAS) (Berginski et al., 2011; Berginski ME and Gomez SM., 2013) picks up adhesions on mock and si*DOCK5* transfected cells. Scale bar is 10 µm. (ii) Focal adhesions tracked over time in same cells as in (i). Blue to red indicates adhesions which have been present for less, and longer amounts of time respectively. (d) (e) Cumming estimation plots of Hedge’s g for comparison between (d) assembly and disassembly rates, and (e) assembly phase length and disassembly phase length of adhesions in mock versus si*DOCK5* transfected cells. N shows number of adhesions quantified, data from 4 cells. Raw data is plotted in the top axes, while mean difference is plotted in lower axes as a bootstrap sampling distribution. Mean differences are shown as dots, 95% confidence intervals are shown by the ends of the vertical error bars. Generated using Ho et al., 2019(Ho et al., 2019) website. Effect size legend |0.0-0.2| no effect (blank), |0.2-0.5| small (S), |0.5-0.8| medium (M), |0.8-1.2| large (L), |1.2-2.0| very large (VL). (d) Unpaired Hedge’s g between mock and si*DOCK5* for assembly and disassembly rates are 1.65 (VL) (95% CI 1.33, 1.96), and 1.28 (VL) (95% CI 1.07, 1.50) respectively. Two-sided P values of Mann-Whitney tests are 3.64 x 10^-15^, and 1.04 x 10^-15^ for assembly and disassembly rates respectively. (e) Unpaired Hedge’s g between mock and si*DOCK5* for assembly phase length and disassembly phase length are 0.101 (95% CI -0.199, 0.405), and -0.00986 (95% CI -0.38, 0.155) respectively. Two-sided P values of Mann-Whitney tests are 0.602, and 0.437 for assembly and disassembly phase lengths respectively. N number is individual adhesions from 4 cells, filtered by them being present for a minimum of 5 frames (phase length 5) and mean axial ratio less than 3. Unpaired Mann-Whitney test, P < 0.001.

To determine if DOCK5 has a role in FA dynamics in LM2 cells which may explain the polarity defects, we used Total Internal Reflection Fluorescence (TIRF) microscopy to visualise the recruitment of Talin-mApple to FAs in migrating mock and DOCK5 depleted cells. In migrating mock transfected LM2 cells, small Talin-positive FAs were established 0-4 µm from the leading edge in the lamellipodia of migrating cells (Fig. 8b and ci, adhesions coloured by FAAS software; Methods, Supplementary Movie 1). These FAs then enlarged/matured and turned over in the same region of attachment to the ECM, and FAs ere not stable in the lamella and/or cell body as the cell moved over this region (Fig. 8cii – dynamics, Supplementary Movie 1).

While migrating mock cells established, and maintained, a polarised leading edge which is the site of both FA formation and turnover (Fig. 8ci and Supplementary Movie 1), the changes in FA dynamics in DOCK5 depleted cells coincided with a failure of these cells to maintain polarity (Fig. 8cii and Supplementary Movie 2). Once established, the leading edge in randomly migrating control cells was maintained for approximately 120 min. In DOCK5 depleted cells there was rarely only one leading edge at any given time but rather many protrusions lasting approximately 40 minutes on average (Supplementary Movie 2). Thus DOCK5 cells were capable of initial establishing polarised protrusions, but the protrusions were not stable. These data demonstrate that DOCK5 depleted cells can initiate the formation of a polarised leading edge but fail to maintain it.

We quantified the assembly and disassembly dynamics of individual FAs in mock and DOCK5 depleted cells. We used a Cumming estimation plot to demonstrate the mean difference in the assembly and disassembly rates and phase lengths. Effect sizes based on Cohen’s initial suggestions (Cohen, 1988; Sawilowsky, 2009) indicate that values of 0.2 to 0.5 are small, 0.5 to 0.8 are medium, 0.8 to 1.2 are large, and 1.2 to 2.0 are very large effect sizes. In DOCK5 depleted cells, FAs have significantly accelerated assembly and disassembly rates compared to control FAs (Fig. 8d, orange unpaired Hedges’ g of 1.65; red unpaired Hedges’ g of 1.28). But disassembly phase lengths were largely unaffected (Fig. 8e, Hedge’s g of 0.101 and -0.0986 for assembly and disassembly phase lengths respectively). Despite accelerated disassembly rates, FAs in DOCK5 depleted cells often fail to completely turnover (Fig. 8b, c) and DOCK5 depleted cells shift their mass over incompletely turned over FA structures as they migrate such that FAs appear to be near the nucleus (Fig. 8cii, Supplementary Movie 2).

The most parsimonious explanation for these observations is that in the absence of DOCK5, nascent FAs can be rapidly stabilised, but are also highly unstable and exhibit fast shrinkage rates, potentially because actin-mediated force maturation is also impaired in DOCK5 depleted cells (Case and Waterman, 2015). Meaning, FAs in DOCK5 depleted cells are poorly coupled to RHOA-dependent actomyosin activity and do not mature properly. However, the absence of actomyosin engagement at FAs in DOCK5 depleted cells also means that turnover of existing FAS structures is also impaired, and thus FAs are “left behind” as the cell moves over them (Fig. 8cii). This model is consistent with the observation that lamellipodia are highly unstable in DOCK5 depleted cells (Supplementary Movie 2).

### CDC42 and RHOA depletion affect FA morphogenesis in metastatic cells

We investigated whether the defects in FA morphogenesis following DOCK5 depletion could be attributed to decreased DOCK5-mediated RAC activation, or whether DOCK5 regulates FA morphogenesis in RAC-independent ways. We first knocked down RAC1, RHOA, and CDC42 and measured Talin-mApple dynamics in migrating LM2 cells (Fig. 9a). We also measured FA dynamics following treatment with Uprosertib to determine the effect of AKT signalling on FA morphogenesis.

**Fig. 9.**
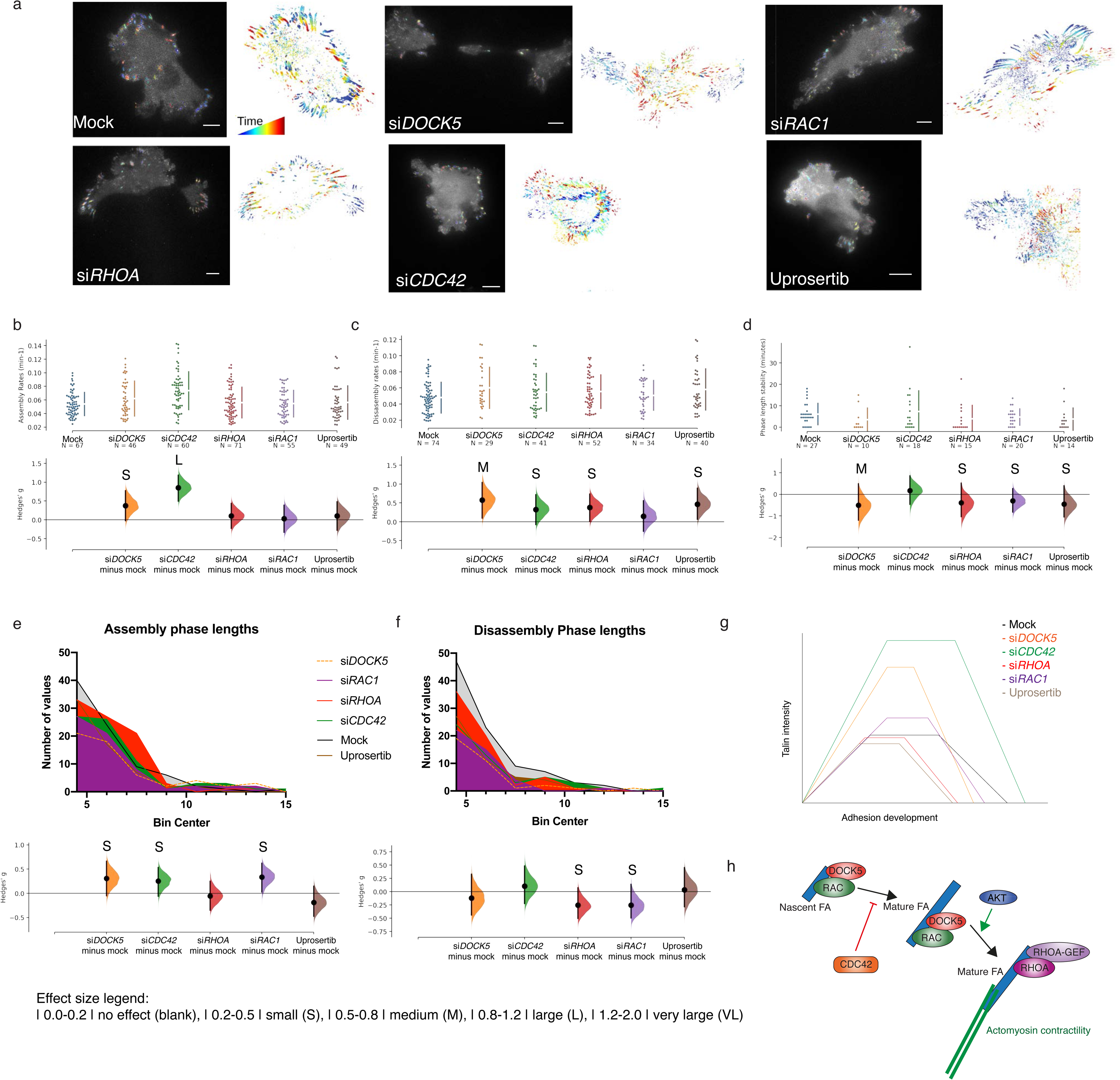
CDC42 and RHOA depletion affect FA morphogenesis in metastatic cells. (a) Representative images of LM2-CAAXGFP cells transfected with Talin mApple and si*DOCK5*, si*CDC42*, si*RHOA*, si*RAC1*, mock, and treated with the AKT inhibitor Uprosertib. Left and right panels per condition show FAAS picking up the adhesions and adhesions tracked over time for the same cell. (b) (c) (d) Cumming estimation plots of Hedge’s g for comparison between (b) assembly and (c) disassembly rates, as well as (d) stability phase length of adhesions in mock versus si*DOCK5,* si*CDC42*, si*RHOA*, si*RAC1* transfected cells, and Uprosertib treated cells. n shows number of adhesions. Data is taken from 3 cells per condition. Raw data is plotted in the upper axes, while mean difference is plotted in lower axes as a bootstrap sampling distribution. Mean differences are shown as dots, 95% confidence intervals are shown by the ends of the vertical error bars. Generated using Ho et al., 2019(Ho et al., 2019) website. Effect size legend |0.0-0.2| no effect (blank), |0.2-0.5| small (S), |0.5-0.8| medium (M), |0.8-1.2| large (L), |1.2-2.0| very large (VL). (b) Unpaired Hedge’s g for assembly rates between mock and conditions: si*DOCK5*: 0.375 (S) (95% CI -0.0214, 0.776); si*CDC42*: 0.851 (L) (95% CI -0.499, 1.19); si*RHOA*: 0.106 (95% CI -0.229, 0.436); si*RAC1*: 0.0311 (95% CI -0.332, 0.387); Uprosertib: 0.105 (95% CI -0.276, 0.479). Two-sided P values of Mann-Whitney tests are 0.242, 2.090 x 10^-5^, 0.922, 0.963, 0.808 respectively. (c) Unpaired Hedge’s g for disassembly rates between mock and conditions: si*DOCK5*: 0.574 (M) (95% CI 0.0979, 1.04); si*CDC42*: 0.321 (S) (95% CI -0.0767, 0.715); si*RHOA*: 0.376 (S) (95% CI 0.0138, 0.732); si*RAC1*: 0.145 (95% CI -0.252, 0.558); Uprosertib: 0.467 (S) (95% CI 0.0641, 0.887). Two-sided P values of Mann-Whitney tests are 0.0363, 0.1840, 0.0580, 0.4030, 0.0652 respectively. (d) Unpaired Hedge’s g for stability phase lengths between mock and conditions: si*DOCK5*: -0.507 (M) (95% CI -1.20, 0.494); si*CDC42*: 0.167 (95% CI -0.464, 0.856); si*RHOA*: - 0.395 (S) (95% CI -1.03, 0.517); si*RAC1*: -0.299 (S) (95% CI -0.827, 0.259); Uprosertib: -0.456 (S) (95% CI -1.04, 0.393). Two-sided P values of Mann-Whitney tests are 0.0602, 0.6390, 0.06080, 0.4590, 0.0476 respectively. (e) Histogram of assembly phase lengths for all conditions (top) and unpaired Hedge’s g for assembly rate phase lengths between mock and conditions (bottom): si*DOCK5*: 0.307 (S) (95% CI -0.0508, 0.662); si*CDC42*: 0.256 (S) (95% CI -0.0605, 0.536); si*RHOA*: -0.0537 (95% CI -0.35, 0.247); si*RAC1*: 0.336 (S) (95% CI 0.0297, 0.621); Uprosertib: -0.187 (95% CI -0.471, 0.148). Two-sided P values of Mann-Whitney tests are 0.140, 0.154, 0.471, 0.237, 0.390 respectively. Mock n = 83, si*DOCK5* n = 56, si*CDC42* n = 74, si*RHOA* n = 83, si*RAC1* n = 67, Uprosertib n = 61. (f) Histograms of disassembly phase lengths (top) and unpaired Hedge’s g for disassembly rate phase lengths between mock and conditions (bottom): si*DOCK5*: -0.12 (95% CI -0.439, 0.39); si*CDC42*: 0.106 (95% CI -0.224, 0.483); si*RHOA*: -0.255 (S) (95% CI -0.505, 0.0726); si*RAC1*: -0.251 (S) (95% CI -0.494, 0.14); Uprosertib: 0.037 (95% CI - 0.285, 0.456). Two-sided P values of Mann-Whitney tests are 0.550, 0.633, 0.270, 0.435, 0.818 respectively. Mock n = 92, si*DOCK5* n = 35, si*CDC42* n = 50, si*RHOA* n = 64, si*RAC1* n = 42, Uprosertib n = 50. (g) Summary of results and how each siRNA and treatment is affecting adhesion life cycle. Not to scale. (h) Cartoon summarising our data where CDC42 inhibits FA growth and turnover, RAC1 promotes FA growth, and RHOA and AKT promote engagement of FAs with actomyosin leading to FA maturation (stability) as well as normal turnover.

Depleting CDC42 significantly accelerated assembly rates (Hedges’ g: 0.851) in a similar manner to DOCK5 depletion, and large FAs formed rapidly (Fig. 9b) (Supplementary Movie 3). However, all other aspects of FA morphogenesis including disassembly rates (Fig. 9c), the length of assembly (Fig. 9e), stability (Fig. 9d), and disassembly phase length (Fig. 9f) were largely unaffected.

In contrast, RAC1 depletion increased the length of the assembly phase (Fig. 9e); consistent with the role of RAC1 in promoting FA assembly (Pasapera et al., 2015) (Supplementary Movie 4). RHOA depletion resulted in a significant increase in FA disassembly rates (Fig. 9c), in line with the idea that in the absence of RHOA-mediated actomyosin engagement, FAs fail to mature (Supplementary Movie 5). Uprosertib treatment resulted in similar effects as RHOA inhibition, as FAs had high disassembly rates, indicative of unstable adhesions (Fig. 9c) (Supplementary Movie 6). However the effect on FA morphogenesis based on all other metrics following depletion of RAC1 and RHOA, or following Uprosertib treatment, was relatively mild.

Taken together these data suggest that CDC42 inhibits FA growth and turnover, RAC1 promotes FA growth, while RHOA and AKT promote engagement of FAs with actomyosin leading to FA maturation (stability) as well as normal turnover (Figure 9h). DOCK5 depletion appears to have much broader effects on FA morphogenesis than any single Rho GTPase alone suggesting it acts upstream of all Rho GTPases – either directly as a GEF or indirectly - in the control of FA growth, maturation and turnover. Further, AKT inhibition closely mimics that of RHOA, suggesting AKT functions in the late stages of FA morphogenesis.

### DOCK5 and CDC42 stabilise polarity

We next sought to explore the idea that DOCK5 acts to modulate a CDC42-GSK3β axis that stabilises MTs at the leading edge of protrusive cells to reinforce the establishment of polarity. Cells were plated on fibronectin (FN) and fixed after 3 hours. After 3 hrs, spreading LM2 cells establish a single leading edge adhered to the matrix by stable FAs, and thus have an asymmetric morphology (Fig. 10). In regions with large FAs, MTs were typically extended into the protrusions (Fig 10 zoomed insert, arrows) supporting the idea that MTs are captured at the cortex in regions with active FA morphogenesis.

**Fig. 10.**
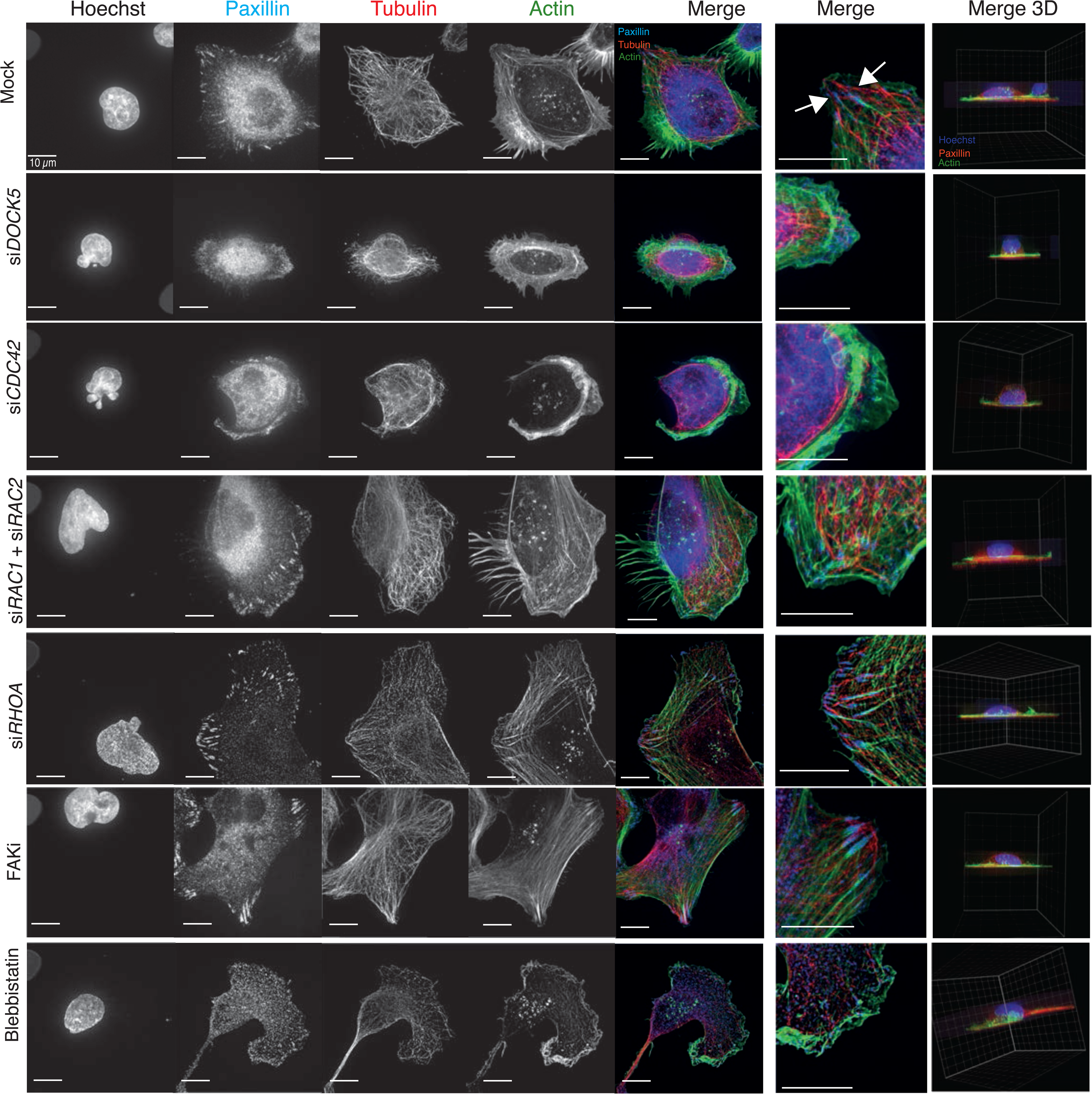
DOCK5 and CDC42 stabilise polarity. Images taken on SoRA for mock, si*DOCK5*, si*CDC42,* si*RAC1+* si*RAC2,* si*RHOA,* and treated with FAKi, and Blebbistatin. Stained for DNA, Paxillin, α-tubulin, F-actin. Merge 2D projection and zoomed section: Paxillin (blue), α-tubulin (red), F-actin (green). Merge 3D projection: Paxillin (red), DNA (blue), F-actin (green). Scale bar = 10µm.

After 3 hrs of spreading, DOCK5 depleted cells established multiple smaller protrusions, leading to asymmetric shapes (Fig. 10). Paxillin-positive FAs in DOCK5 depleted cells were very small and appeared diffusely spread in the leading edge (Fig. 10 zoomed insert). Consistent with previous reports demonstrating a role of DOCK5 in promoting MT stability (Guimbal et al., 2019; Ogawa et al., 2014), there were dramatically fewer MT filaments in the periphery of DOCK5 depleted cells, and MTs appear thin and poorly organized compared to those of mock transfected cells. In DOCK5 depleted cells MTs appear organised largely as bundles around the nucleus. MT bundles are often hallmarks of stabilised, old MTs that exhibit low instability and are not captured at the cortex (Gundersen, 2002). Thus we conclude DOCK5 is essential for MT capture at the cortex in protrusions.

CDC42 depletion led to a partially similar phenotype as that of DOCK5, as cells formed expanded lamellipodia that nearly encompassed the entire circumference of the cells, and adopted a rounded, symmetric shape (Fig. 10). However, Paxillin-positive FAs in CDC42 depleted cells appeared globular and poorly formed – consistent with live cell imaging data suggesting CDC42 inhibits FA assembly (Fig 10). In contrast, cells depleted of RHOA, or RAC1+RAC2 all formed polarised lamellipodia with multiple large FAs (Fig. 10), hallmarks of defective FA maturation and/or turnover, but not in MT organization or establishment of polarity. Treatment of cells with either a focal adhesion kinase inhibitor (FAKi), which prevents focal adhesion turnover; or Blebbistatin, a contractility inhibitor, did not inhibit the formation of polarised lammelipodia, although no FAs formed in Blebbistatin treated cells (Fig. 10). Thus we conclude DOCK5 promotes the stability of polarised leading edge by acting at FAs to promote CDC42-mediated MT capture at the cortex.

### DOCK5 promotes spreading following MT depolymerisation

DOCK5 has been shown to promote MT stability downstream of GSK3β phosphorylation (Guimbal et al., 2019; Ogawa et al., 2014). To investigate the relationship between MTs and DOCK5 at the leading edge, we performed assays to analyse cell morphology following washout of the MT-depolymerization agent nocodazole (NZ) in media containing either: NZ, the MT-stabilising agent taxol (TX) or DMSO. Following 4 hrs of NZ both mock and DOCK5 depleted cells showed decreased spread area, adopted a rounded morphology and actin intensity at the membrane was significantly decreased (Fig. 11a, b, c). However, while in control cells MT staining was expectedly diffuse, in DOCK5 depleted cells tubulin was localised into large high-intensity “bundles” which was reflected by significant increases in the SER Ridge Feature (texture feature) (Fig. 11d). These data show that MTs are differentially affected by MT depolymerising agents in control vs DOCK5 depleted cells, and we propose that depletion of DOCK5 leads to MT bundling and stability because MTs are not captured at the cell cortex (Gundersen, 2002).

**Fig. 11.**
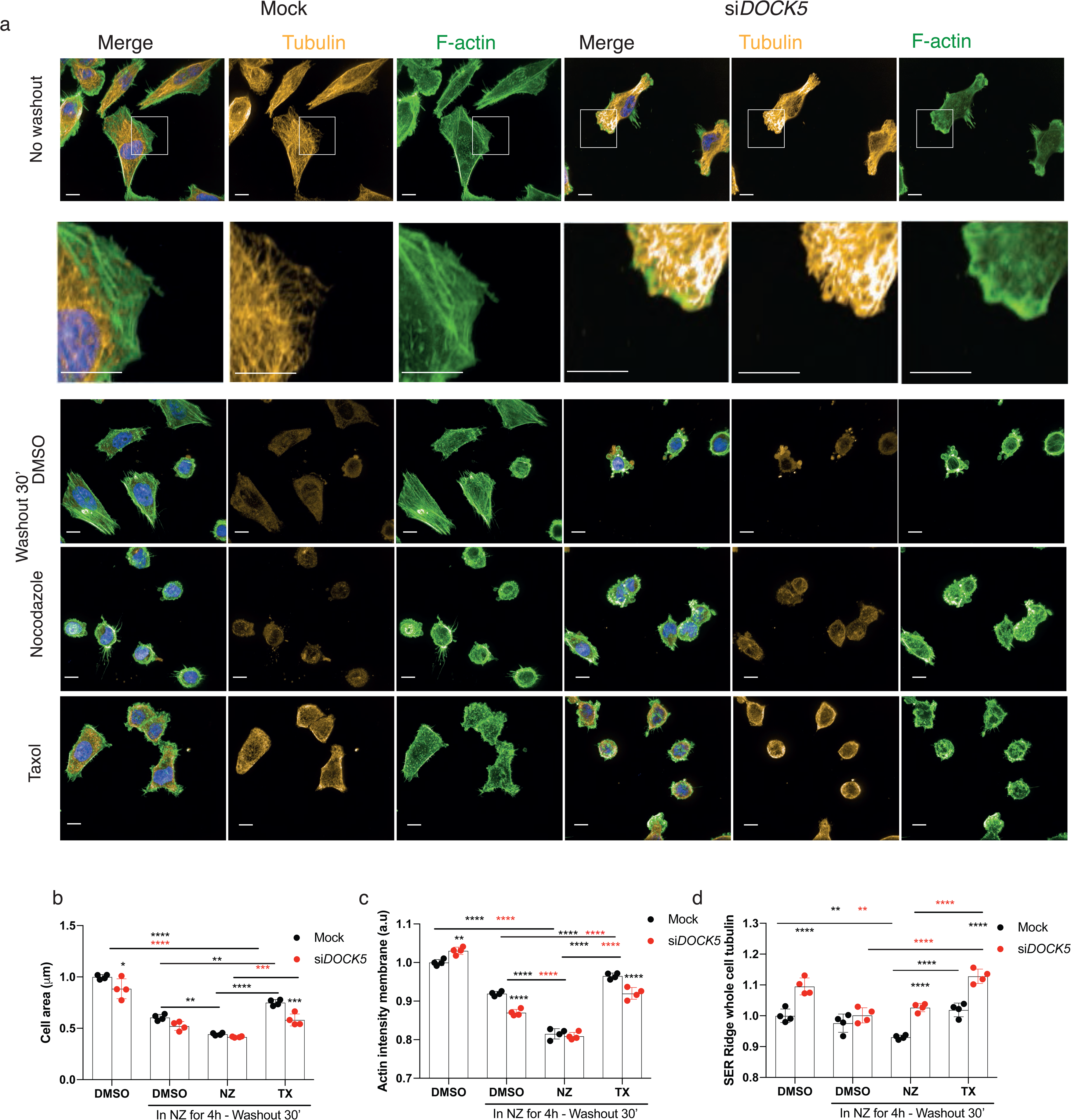
DOCK5 promotes spreading following MT depolymerisation. (a) Immunofluorescence images (maximum projections) of mock or si*DOCK5* transfected cells treated just with DMSO or treated with nocodazole (NZ) for 4 hours and then washed out in either media containing DMSO, NZ, or taxol (TX). Stained for DNA (blue), tubulin (yellow), and F-actin (green). Scale bar is 10 µm. Imaged at 60x with water objective (NA = 1.2). Images shown with same intensity settings. White box represents zoomed in sections below. (b-d) Quantifications of (a) showing (b) cell area, (c) actin intensity at the membrane, and (d) whole cell SER Ridge tubulin (texture). Mean per well ± sd are shown (n > 3 wells, 1000 cells/well, 2 ways ANOVA, Tukey’s multiple comparison’s test * P < 0.05, ** P < 0.01, *** P < 0.001, **** P < 0.0001). Significance between Mock and si*DOCK5* for each condition shown with no bars. Significance between mock NZ (black) and si*DOCK5* NZ (red) compared to other treatments for same transfection.

Thirty minutes after washout in DMSO containing media, control cells formed abundant filopodial protrusions and increased in cell area (Fig. 11a, b, p = 0.013) likely as a result of MT outgrowth and RAC1 promoting actin polymerisation(Waterman-Storer et al., 1999; Wittmann et al., 2003). Indeed, actin intensity at the membrane significantly increased in DMSO media washout denoting actin polymerisation (Fig. 11c). DOCK5 depleted cells also spread and increased in cell area under DMSO containing media washout (Fig. 11b, p = 0.013 for mock, p = 0.0894 for si*DOCK5*) but had lowered levels of actin at the membrane (Fig. 11c) – supporting the idea that DOCK5 is required for the stability of actin-based protrusions. DOCK5 depleted cells did not display polarised fronts for any condition supporting our hypothesis that DOCK5 is required for maintenance of polarity. Notably washout of NZ in DOCK5 depleted cells affected MT organisation in an identical manner as control – effectively rescuing the effects of DOCK5 depletion on MT organisation following NZ treatment. Therefore it is unlikely DOCK5 regulates polymerisation of MTs during spreading or migration, but rather regulates MT capture.

Washout in TX containing media resulted in a significant increase in cell area for both mock and DOCK5 depleted cells compared to washout in DMSO alone, effectively accelerating the rate of spreading, though the ability of TX to rescue spreading was blunted in DOCK5 depleted cells (Fig. 11a, b, p = 0.005 for mock, p = 0.9114 for si*DOCK5*). Accelerated spreading was likely due to increased actin polymerisation, as both mock and DOCK5 depleted cells in TX had increased levels of actin at the membrane compared to DMSO washout alone, and were at levels comparable to controls. Strikingly however, TX resulted in significant increase of MT bundles in DOCK5 treated cells, that were even higher than the DMSO treated DOCK5 depleted cells (Fig. 11d). We propose that the formation of MT bundles in TX washout is due to accelerated MT polymerisation due to TX, but a failure to capture MT plus ends at the leading edge.

## Discussion

In both yeast and mammalian cells the recruitment and activation of CDC42 at the membrane serves to seed polarised cortical structures that are enriched complexes and cytoskeletal elements that would otherwise be diffusely or asymmetrically organised in cells (Chiou et al., 2017). During the formation of the bud in *S. cerevisiae*, localised CDC42 activation marks the site of bud assembly at one site on the membrane (Johnson et al., 2011; Richman et al., 2002). A key aspect of bud morphogenesis are positive feedback loops initiated by CDC42 which increase the local concentration of GTP-bound and active CDC42 (Kozubowski et al., 2008). One effect of these loops is that one bud site outcompetes all other potential sites for the recruitment of bud assembly in a “winner takes all” fashion (Howell et al., 2012; Wu et al., 2015). Indeed in yeast mutants where the reinforcement, but not necessarily the establishment, of polarity is disrupted, multiple bud sites form (Freisinger et al., 2013; Wu et al., 2015).

In the case of migrating mammalian cells, the establishment of polarity occurs with the coincident reorganisation of the actomyosin cytoskeleton to generate protrusions such as lamella and lamellipodia, and the engagement of the ECM by integrins to form FAs. Classical studies have shown that CDC42 regulates microtubule stability and establishment of polarity via PAR6 and PKCζ mediated phosphorylation of GSK3β at the leading edge of migrating cells in a manner that is integrin dependent (Etienne-Manneville, 2004; Etienne-Manneville and Hall, 2003b, 2003a). Here we propose that DOCK5 acts at FAs in breast cancer cells to establish positive feedback loops which maintain polarity in the presence of stable integrin-ECM attachments and cellular tension. By reinforcing polarity in sub-cellular regions with stable FA attachments, DOCK5 promotes processive migration and 3D invasion (Fig. 12). We and others have previously observed that DOCK5 is recruited to FAs (Frank et al., 2017; Müller et al., 2020), and here we show that DOCK5 acts to couple FA morphogenesis and polarity by coordinating multiple GTPase activities. Firstly, DOCK5 acts as a GEF for RAC-type GTPase to promote FA assembly and ultimately maturation/turnover. In the absence of DOCK5, FAs do not increase in size, and the turnover of existing FAs is disrupted (Fig. 8) likely because of a failure of DOCK5-depleted FAs to engage actin at the leading edge (Case and Waterman, 2015; Hall, 1998; Nobes and Hall, 1995). Failure to mature FAs and/or engage actomyosin likely results in decreases in RHOA activation (Fig. 12). Secondly, by recruiting and activating AKT to FAs near the leading edge, DOCK5 stabilises GSK3β, which is targeted by CDC42-PKCζ for phosphorylation, leading to the maintenance of polarity. In cells depleted of DOCK5, CDC42 activation at the membrane, the initial establishment of polarity, is initially unaffected. But either C21 treatment or lack of DOCK5 reduces the levels of GSK3β that can be phosphorylated by CDC42-PKCζ, leading to a failure of cells to maintain polarity. CDC42 activation (which is initially independent of DOCK5) at another subcellular site can lead to formation of a new unstable leading edge in other regions of the cells. Indeed, a characteristic of DOCK5 depleted cells is not the ability to form protrusions, but rather to maintain a stable polarised protrusion. As such this phenotype is highly reminiscent of *rdi1* and *bem1/2* yeast mutants which form numerous unstable buds (Freisinger et al., 2013; Kozubowski et al., 2008; Wu et al., 2015).

**Fig. 12.**
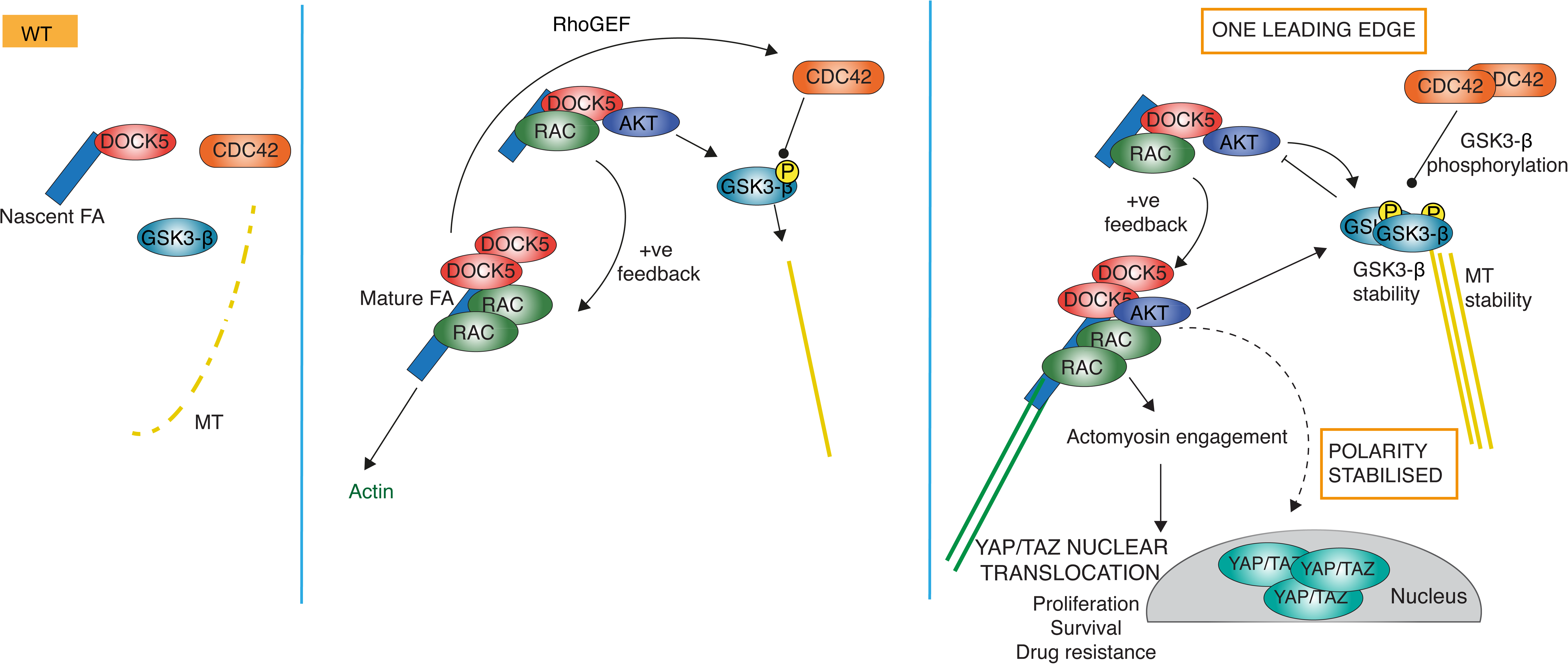
DOCK5 acts at FAs in breast cancer cells to establish positive feedback loops which maintain polarity in the presence of stable integrin- ECM attachments and cellular tension. Schematic of proposed model where in WT conditions DOCK5 gets recruited to FAs and acts as GEF for RAC-type GTPase to promote FA assembly and turnover/maturation in a positive feedback loop. At the same time DOCK5 recruits and activates AKT to FAs near the leading edge stabilising GSK3β, which is targeted by CDC42-PKCζ for phosphorylation, leading to the maintenance of polarity. Here DOCK5 increases the levels of AKT and thus promotes further GSK3β stability and phosphorylation in another positive feedback loop. Further, actomyosin engagement by FA results in high RHOA and consequently high nuclear YAP/TAZ. Finally, DOCK5 could potentially mature FAs to recruit CDC42 GEFs which can further amplify CDC42 activation at the leading edge.

Critically, by promoting FA assembly we propose DOCK5 initiates positive feedback loops which stabilise protrusions during migration. DOCK5 activity at FAs can promote FA assembly leading to the recruitment of more DOCK5 complexes, actomyosin engagement and turnover (Müller et al., 2020). DOCK5 complexes can also recruit and activate AKT which promotes GSK3β stability and phosphorylation (Fig. 12). Potentially DOCK5 mediated FA maturation can also lead to the recruitment of CDC42 GEFs which can further amplify CDC42 activation at the leading edge. Indeed depletion of CDC42 GEFs FGD5 and PLEKHG3 (Ando et al., 2013; Nguyen et al., 2016) results in similar morphological changes as DOCK5 depletion (Fig. 5). FGD5 has been previously shown to regulate polarity downstream of Rap1 GTPase (Ando et al., 2013), and thus could serve an orthologous function to yeast CDC24 RhoGEF which is activated by Rap1 signalling to promote CDC42 positive feedback at the bud scar (Kozubowski et al., 2008).

DOCK5 has been previously described to recruit and activate AKT and promote GSK3β phosphorylation through both RAC-dependent and independent ways (Guimbal et al., 2019; Ogawa et al., 2014). In particular, DOCK5’s C-terminus has been shown to recruit NCK/AKT complexes to promote GSK3β phosphorylation in ways that are independent of catalytic activation of RAC-type GTPase through its DHR2/GEF domain (Guimbal et al., 2019; Ogawa et al., 2014). But our observation that the DOCK5 GEF inhibitor C21 mimics the effect of both DOCK5 siRNA and AKT inhibition by Uprosertib on GSK3β localisation at the membrane and YAP/TAZ translocation strongly suggests that DOCK5 mediated activation of RAC-type GTPases via its DH2/GEF is required for GSK3β phosphorylation and localisation at the leading edge. One model which resolves these previously contradictory observations is that activated DOCK5 exists in a conformation that recruits NCK and AKT to its C-terminus independent of it GEF domain; but also functions as a GEF to promote RAC-mediated FA assembly – which recruits more DOCK5/NCK/AKT to FAs in a positive feedback loop. Thus inhibition of RAC by C21 breaks this loop and lowers AKT activation. Our observation that AKT inhibition by Uprosertib mimics different aspects of DOCK5 depletion on FA morphogenesis in fact suggests that AKT activity itself promotes FA morphogenesis and is involved in this positive feedback loop (Fig. 12).

We and others have previously demonstrated that YAP/TAZ nuclear translocation is dependent on cell shape and geometry (Dupont et al., 2011; Pascual-Vargas et al., 2017; Piccolo et al., 2014; Sero and Bakal, 2017; Totaro et al., 2018). Our data suggest that decreases in YAP/TAZ nuclear translocation in DOCK5 depleted cells are due to the inability of DOCK5 depleted cells to maintain a polarised, adherent morphology leading to low YAP/TAZ. Mechanistically this can be explained by the inability of cells to assemble stable FAs (Fig. 8). The absence of fully mature FA leads to both defects in assembly of complexes at FA which promote YAP/TAZ activation – i.e. via signalling pathways such as PAK2 (Sero and Bakal, 2017) or FAK-PI3K (Kim and Gumbiner, 2015) and/or the failure of FAs in DOCK5 depleted cells to engage actin and upregulate actomyosin contractility (Etienne-Manneville, 2008; Ridley et al., 2003) which promotes YAP activation (Calvo et al., 2013). This model is perhaps best supported by the finding that RHOA depletion, and a failure to mature FAs leads to significant decreases in YAP/TAZ that are comparable to complete loss of polarity following CDC42 depletion (Fig 1a, 9c, 10).

It is tempting to speculate that sustained YAP/TAZ nuclear translocation, and resulting capabilities such as drug resistance, may be a consequence of cancer cells adopting stable polarised migratory forms. In the case of LM2 cells, the idea that processive migratory forms driven by DOCK5 activity leads to shape changes which then drive YAP/TAZ activity and cancer relevant phenotypes such as drug resistance, is further supported by the finding that across genetic backgrounds (different RNAi perturbations) shape is predictive of YAP/TAZ activity (Fig. 4e, f). This model would be in contrast to the classical idea that drug resistance and migration are coincident events, and/or that YAP/TAZ itself drives migration. Indeed, studies in melanoma cells have shown that YAP/TAZ upregulation which promotes drug resistance is caused by shape changes (Fisher et al., 2017; Kim et al., 2016; Lin et al., 2015a; Lionarons et al., 2019). Moreover, a number of recent studies have shown that EMT in breast cancer, which is characterized by morphological changes and increased invasive capabilities, also leads to drug resistant cells (Dongre and Weinberg, 2019). Thus shape changes may be causal to many aspects of TNBC, and suggests that therapeutic intervention could occur by preventing cells from altering their shape.

## Materials and Methods

### Cell culture

LM2s (subpopulation 4172 from MDA-MB-231) were obtained from Joan Massagué (Sloan Kettering Institute, New York) (Minn et al., 2005), and stably express a triple-fusion protein reporter construct encoding herpes simplex virus thymidine kinase 1, green fluorescent protein (GFP) and firefly luciferase (Minn et al., 2005). MDA-MB-231 cells were obtained from Janine Erler (University of Copenhagen, Denmark), while SUM159 and hs578t were a gift from Rachel Natrajan (Institute of Cancer Research, London). Cells were grown in Dulbecco’s Modified Eagle Medium (DMEM) supplemented with 10% heat-inactivated fetal bovine serum (FBS) and 1% penicillin/streptomycin in T75 falcons, at 37**°**C and supplemented with 5% CO_2_ in humidified incubators. MCF10A were obtained from ATCC and were grown in DMEM/F12 supplemented with 5% horse serum, 10µg/ml insulin, 20ng/ml epidermal growth factor, 100ng/ml cholera toxin, 500ng/ml hydrocortisone, and 100mg/ml penicillin/streptomyocin in T75 falcons. T47D were obtained from from Rachel Natrajan (ICR), and were grown in Roswell Park Memorial Institute (RPM1)-1640 culture medium (Gibco) supplemented with 10% heat-inactivated fetal bovine serum (FBS) and 1% penicillin/streptomycin in T75 falcons.

### siRNA transfections

All siRNAs were made up to a stock of 20µM. Briefly, for 384 well plates, siRNA reverse transfections were performed by adding 40nL of siRNA to each well followed by 5µl of Opti-MEM® Reduced Serum Media. 5 minutes later 5µl of mix containing Opti-MEM and RNAimax reagent in a 125:1 ratio were added per well. Plates were spun at 1000 rpm for 1 minute, and incubated at room temperature for 20 minutes to allow siRNA-RNAimax complexes to form. Cells were plated on top in 30µL of DMEM (10% FBS, 1% penicillin/streptomyocin), resulting in a total volume of 40µL per well: 40nL siRNA (final concentration of 20nM), 5µL Opti-MEM, 5µL of Opti-MEM and RNAimax mix (125:1) plus 30µL of cells. For transfections performed on 96 well plates and 6 well plates, 4 and 64 times the amounts used for 384 well plates were used, respectively. When siRNAs were transfected in combination, the same amount of each siRNA was used. siRNAs and sequences used can be found in Table 1.

**Table 1:**
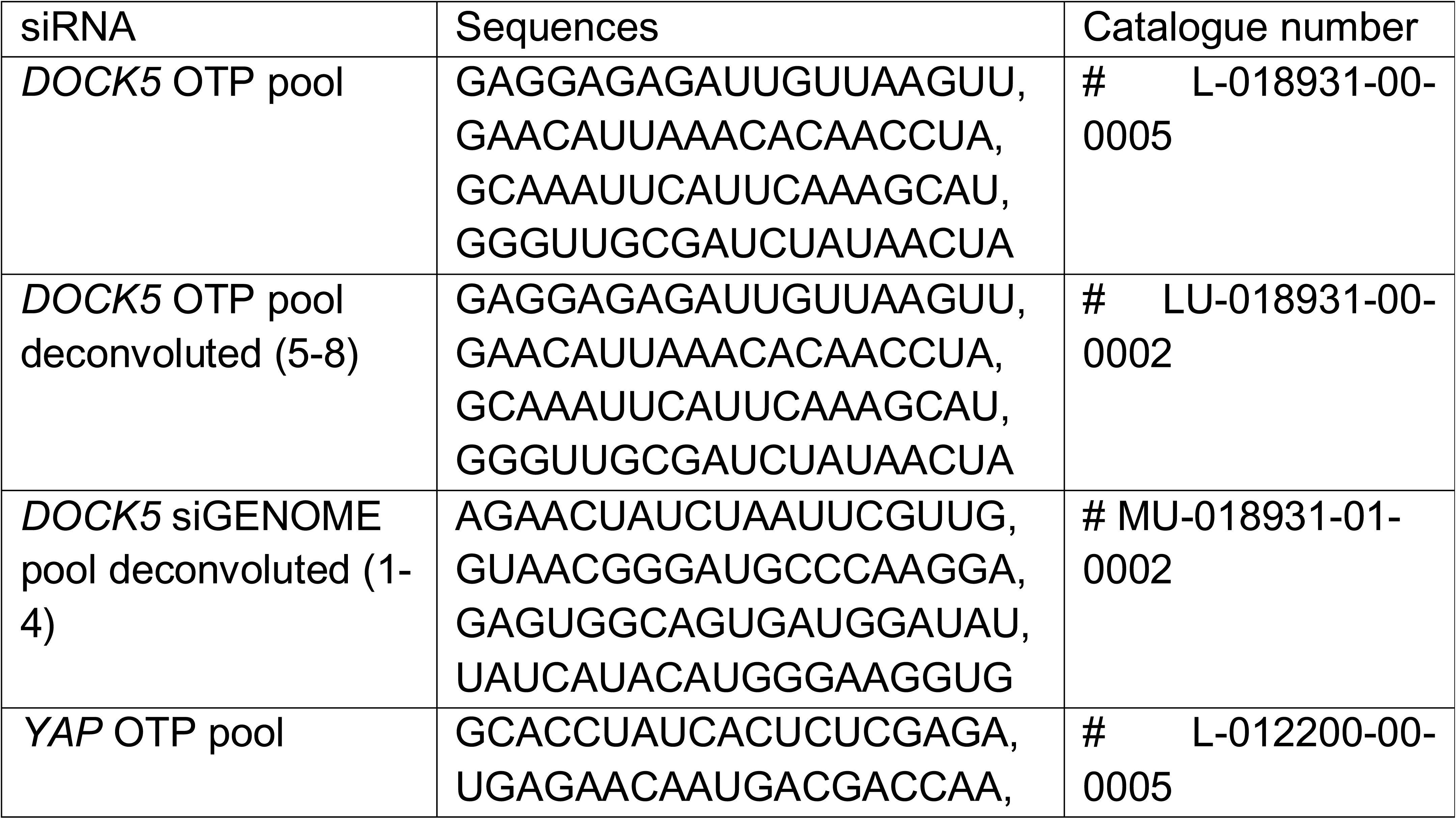
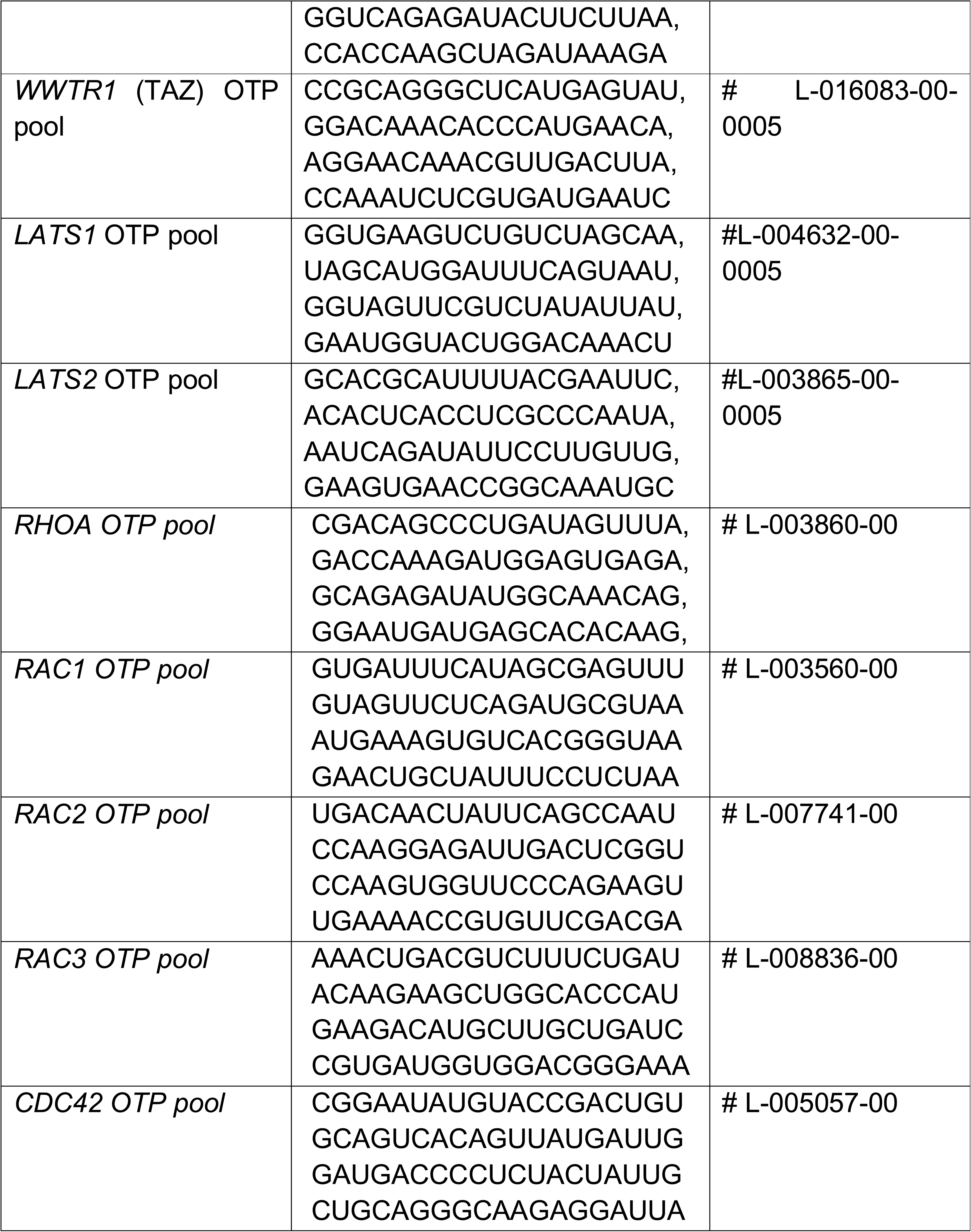
siRNA target sequences

### Plasmid transfections

For transfections with Talin mApple and DOCK5-GFP, Effectene Transfection Reagent (Qiagen) was used according to manufacturer’s protocol. Cells were plated at a density of 1.0 x 10^6^ cells /ml in 1ml in a 6 well plate. Then, 2.56ng of plasmid were combined with 150µl of EC buffer and vortexed followed by addition of 8µl of Enhancer and vortexing. After 10 minutes 5µl of Effectene were added, mix was vortexed, 10 minute later 1ml of DMEM was added and mixture was dispensed dropwise per well. If siRNA was used as well, cells were reverse transfected as usual. Experiments were performed 48 hours later.

### Generation of CAAX-GFP tagged cell line

CAAX-GFP tag was a kind gift from Nicola Ferrari (previously Institute of Cancer Research, currently Astex Pharmaceuticals, UK), who generated it in HEK-293 cells using a second-generation lentiviral vector (plasmid EGFP-CAAX pCSII-IRES2-hygro). Cells were plated at a density of 6.4 x 10^4^ cells /ml in a 6 well plate. Cells were transfected 24 hours later using a 1:1 dilution of lentivirus containing the CAAX-GFP tag solution and DMEM per well according to the manufacturers protocol. Transfected cells were selected with Hygromycin B (Invitrogen, cat # 10687-010) at a final concentration of 500µg/ml (elucidated from carrying out a concentration curve) 72 hours post transfection. CAAX tag did not affect YAP/TAZ ratio or cell morphology.

### Drug treatments

Drugs were usually added on top of media already present. Appropriate amounts of DMSO or H_2_O depending on what the drug was dissolved in were added to mock transfected cells. A list of drugs used can be found in Table 2. Drugs were added for a period of 4 hours, unless other stated, so as to eliminate any cell cycle effects.

**Table 2:** Drugs added.

| Drug | Inhibitor type/target | Company and catalogue number | Final concentration. |
| --- | --- | --- | --- |
| PF-573228 | FAK inhibitor | Tocris Biosciences, #3239; CAS 869288-64-2 | 2µM |
| C21 | DOCK5 inhibitor | Alfa Aesar, #H60031.MD; CAS 54129-15-6 | 100µM |
| Blebbistatin | Myosin II light chain | Sigma, #B0560; CAS 856925-71-8 | 10µM |
| Binimetinib | MEK inhibitor |  | 10 µM |
| Uprosertib (GSK2141795) | AKT inhibitor |  | 2µM |
| Taxol (Paclitaxel) | Microtubule stabiliser | Sigma, #T7191 | 10µM |
| Nocodazole | Microtubule depolymeriser | Sigma, #M1404 | 10µM |

For the nocodazole and taxol experiment, cells were mock and si*DOCK5* reverse transfected as described in RNAi screening methodology for 48 hours. Then, cells were trypsinised and plated at a density of 1500 cells/well and allowed to settle for 20 hours. Microtubule depolymerisation was induced by nocodazole at a final concentration of 10µM for 4 hours. Washout was performed for 30 minutes in either DMSO, 10µM nocodazole, or 10µM taxol – containing DMEM as indicated in Fig 11.

### RNAi screening methodology

40nl (0.08pmol) of siRNA from an ONTARGET*Plus* human siRNA RhoGEF/GAP and Rho GTPase library (stock concentration of 20µM, Dharmacon) were arrayed using the acoustic liquid handler Echo 550 prior to transfection and kept at -80**°**C. All siRNAs were arrayed in duplicate per plate and each plate was arrayed in duplicate resulting in a total of 4 plates per library. Pre-stamped siRNA plates were thawed at room temperature for 30 minutes prior to use. 5µl of Opti-MEM® Reduced Serum Media were then added per well followed by 5µl of RNAiMax reagent (Invitrogen) and Opti-MEM® mix in a 1:125 ratio. Plates were spun down at 1000 rpm for 1 min and incubated at room temperature for 20 mins. Cells (LM2 and MDA-MB-231) were then seeded at a concentration of 33000 cells/ml (1000 cells per well) in 30µl DMEM supplemented with 12.8% FBS, 1.28% penicillin/streptomyocin (10% FBS and 1% penicillin/streptomyocin final) to account for the 10 µl of mix already present in the well, resulting in a total volume of 40µl per well. Automated handling of plates on the Cell::Explorer robot station was carried out using the PlateWorks software (Perkin Elmer). Plates were fixed 48 hours later with 8% warm formaldehyde in PBS (final concentration 4%) for 15 minutes and permeabilised with 0.1% Triton-X-100 in PBS for 10 minutes. Cells were then blocked in 2% BSA in PBS for 2 hours. All antibody incubations were performed in solution containing 0.5% BSA, 0.01% Triton-X-100 dissolved in PBS in a 1:1000 ratio, with the first primary antibody being added overnight at 4**°**C and the rest being incubated at room temperature for 1.5 hours. Plates were washed 3x with PBS between every step using a Microplate Washer (BioTek). Antibodies were added sequentially to prevent cross reactivity. Antibodies used were: mouse YAP/TAZ (Santa Cruz Biotechnologies, cat # sc-101199), rat α - tubulin (Bio-Rad, cat # MCA77G), AlexaFluor 647 goat anti-mouse (Life Technologies, cat # A21235), AlexaFluor 647 goat anti-rat (Life Technologies, cat # A11077), phalloidin (Invitrogen, cat # A12379). Finally Hoechst (Sigma Aldrich, cat # 33258) was added for 15 minutes, and after washing, cells were left in PBS and plates were sealed. Cells were imaged using an automated Opera Quadrupule Enhanced High Sensitivity (QEHS) spinning-disk confocal microscope (Perkin Elmer) with 20x air lens. 28 fields of view were imaged per well. Cells were plated at increasing densities on columns 1, 2, 23 and 24 to be able to account for density dependent effects. SUM159 and hs578t were plated at a density of 67000 cells/ml (2000 cells/well) and 50000 cells/ml (1500 cells/well) respectively.

Images were processed and analysed using Columbus 2.6.0. Software Platform (PerkinElmer). Briefly, single cells were identified using the nuclear Hoechst signal. α-tubulin intensity was using to segment the cytoplasm. Cells touching the image border were filtered out. Following this, cell morphology and texture features were extracted from all stains as described extensively in Pascual-Vargas et al, *Scientific Data* 2017. In the same script, a cell shape linear classifier was manually trained on visually distinctive cells that represented each category (spindly, large round, triangular, fan, and small and round) and included all the features extracted from the stains except those based on YAP/TAZ. YAP/TAZ ratio was calculated as log ratio of nuclear to ring region (perinuclear region) and was not included in the classifier. The percentage of cells classified into each shape was calculated per well and normalized per plate. Z-scores were then calculated per screen using the average and standard deviation of all mock transfected cells. Each well was assigned a QMS consisting of the Z-score for each shape.

Because we have previously observed that following efficient gene depletion, many cells in a population can appear wild-type (Yin et al., 2013), which can potentially obscure the presence of phenotypes resulting from gene depletion, we identified and filtered normal cells from our data set in the following way. Briefly, we performed principal component analysis (PCA) on single cell dataset comprised of texture and morphological features of populations which were either mock transfected or transfected with siRNAs which scored the highest for enriching for a particular shape. The data were projected into 2D PC space and each data point (cell) colour coded by the shape of each cell (Supplementary Fig. 4a). We defined the region of shape space where different classified cells overlapped in PC space as normal cells, defined as those whose PC coordinates lay within 1 SD of the mean of PC1 and PC2 for the siRNA transfected population (black box, Supplementary Fig. 4b). We trained the new linear classifier on the cells that fell in this space in the mock transfected population (black box, Supplementary Fig. 4a) and defined this population as ‘normal’. Running the classifiers again on the same data, performing the same PCA analysis on this new dataset and removing cells classified as ‘normal’ resulted in cells classified into the different shapes separating better in PC shape space (Supplementary Fig. 4c). Further analyses are described in the results section.

### Quantitative Real-Time PCR (qRTPCR)

RNA was extracted using phenol:chloroform (TRIzol®, ThermoFisher cat # 15596026) and RNAeasy kit (Qiagen cat # 74104) according to the manufacturer’s protocol. RNA was converted to cDNA using a cDNA conversion kit (Applied Biosystems cat # 4387406) according to the manufacturers protocol. qRTPCR was performed on a QuantStudio® Flex Real-Time PCR system, using PCR mastermix SyBR green (Applied Biosystems cat # 4309155) and primers in Table 3.

**Table 3:**
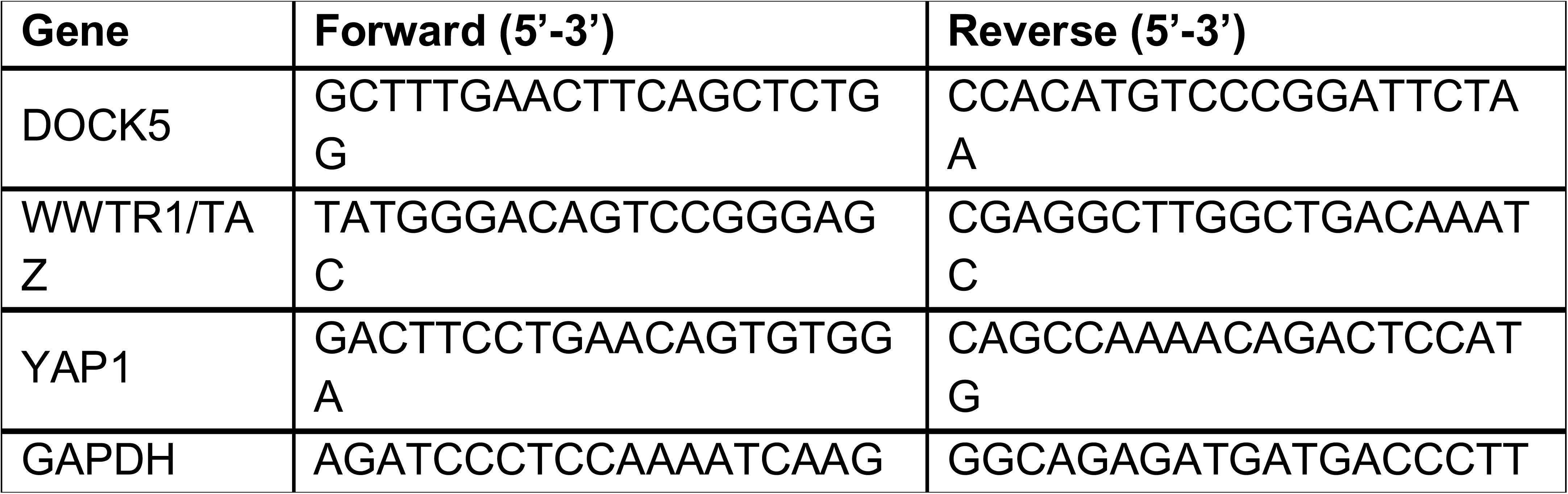
PCR primers used for qRT-PCR.

### 3D collagen invasion assay

We chose rat tail Col-I as our matrix because it is the most abundant type of collagen in mammalian tissues (Brown et al., 2003; Magzoub et al., 2008; Oldberg et al., 2007; Sabeh et al., 2004, 2009), and has been shown to form networks with physiological mechanical properties *in vitro* (Sapudom et al., 2019; Stamov and Pompe, 2012; Wolf et al., 2003).

Cells were reverse transfected in 6 well plates as previously described. 48 hours later cells were trypsinised and resuspended in a solution containing rat-tail collagen I (Corning cat # 354249), ultrafiltered H_2_O (prepared in house according to standard protocols), DMEM 5x and HEPES (1M pH 7.5, prepared in house according to standard protocols). Collagen solutions were prepared on ice and 96 well plates were prechilled prior to use. Collagen solutions were made up as in Table 4 starting with HEPES, followed by DMEM 5X with rat-tail collagen being added last. Neutralisation of collagen was tested with pH strips prior to use to ensure a pH of 7.4. DMEM 5X was made up using 12.5g DMEM powder, 25 ml of 1M HEPES ph 7.5, 5g NaHCO_3_, H_2_O up to 250 ml and was filtered under sterile conditions before aliquoting into 10ml batches and freezing them at -20**°**C. Cells were spun down in 1.5ml eppendorfs at 1000 rpm for 4 minutes, and supernatant was removed. Cell pellets were resuspended carefully so as to not introduce bubbles in 500μL of collagen solution, and 100μL of cell-collagen mix was dispensed per well of a 96 well plate. Once all conditions were dispensed, plates were spun down for 1 minute at 4 **°**C to ensure all cells were at the bottom of the plate. Collagen was allowed to polymerise at 37**°**C for one hour. Then 50 μL of DMEM were added on top of the collagen gels. 150 μL of PBS were added to all outside wells to prevent gel dehydration and edge effects due to dehydration. Cells were fixed 48 hours later with 50 μL/well of 16% PFA containing Hoechst (1:500) for 2 hours. Gels were then washed with PBS and left in PBS prior to sealing. The Opera QEHS spinning-disk confocal microscope (PerkinElmer) with 20x air lens was used to image these plates. A minimum of three Z stacks were taken per well, spaced 30 μm apart with 0 μm referring to the bottom of the plate and subsequent stacks referring to inside the gel. 20 fields were imaged per well per stack, and were all in the centre of the well to prevent edge effects. Columbus software was used to quantify cell number at every stack, using a filtering threshold method to ensure only cells with a certain nuclear and GFP intensity were counted in each stack (Supplementary Fig. 6).

**Table 4:**
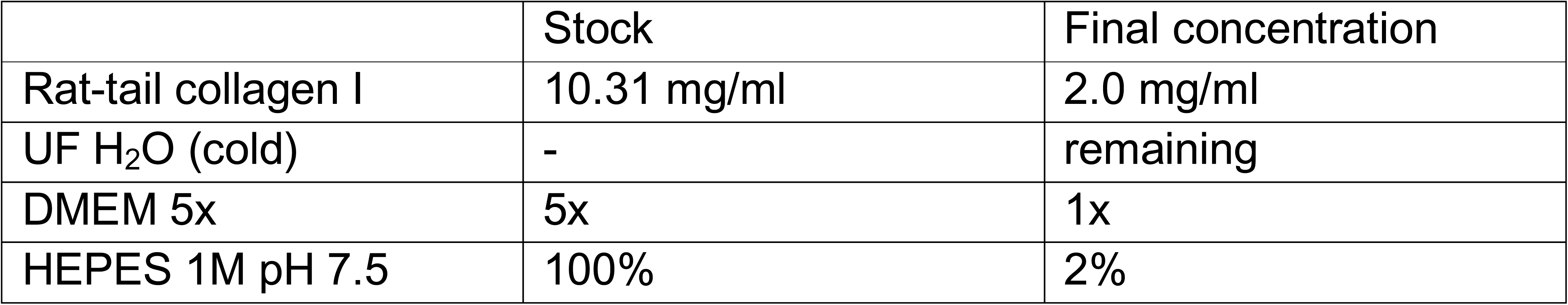
Reagents used for collagen invasion assays.

### Spheroid invasion assay

Cells were reverse transfected in 96 well ultra-low attachment (ULA) plates as described in the siRNA transfections section. Quantities used per well: 20 μl l RNAimax, 20 μl OPTIMEM + 0.16 μl siRNA. Cells were plated at a density of 4.0 x 10^4^ cells/ml in 120 μl, resulting in a final volume of 160μl per well. Due to their mesenchymal nature, LM2 cells required 2% Matrigel in ultra-low attachment plates to form spheroids, therefore 5 hours later a solution of 2.5% Matrigel in 40 μl of DMEM was added per well to promote spheroid formation resulting in a total volume of 200 μl per well. 48 hours later 180 μl of medium was carefully removed from each well so as not to disturb the spheroid, and replaced with 180 μl of 100% Matrigel. Spheroids were imaged prior to adding 100% Matrigel and immediately after using the Celigo Imaging Cytometer (Nexcelom Bioscience). Spheroids were imaged every day for 8 days, following the protocol outlined in (Vinci et al., 2015). Images were processed using ImageJ. Invasion area was determined by drawing a circle to the edge of where cells invaded and subtracting the initial area of the spheroid at day 0. Proliferation was quantified by drawing a circle around the edge of the area of the spheroid where the spheroid is grey, rather than the edge, so as to exclude any invading cells.

### Western blotting

Cells were reverse transfected in 6 well plates at a density of 3.3 x 10^4^ cells /ml in a 6 well plate (same density as screens and all other experiments unless otherwise stated) for 48 hours. Cells were harvested by trypsinising, spinning down in 1.5ml eppendorfs and removing supernatant. Cells were then washed in PBS 3 times. Pellets were then resuspended in 500µl of solution containing a ratio of 3:1 of water to LDS Sample Buffer (4x) (Invitrogen) with a final concentration of 2mM DTT. Samples were then boiled at 100 C for 10 minutes and spun down for 2 minutes at 13 000 rpm. Samples were frozen at -20**°**C. Samples were boiled and spun down again prior to use. 16µl of sample were loaded into 1.0mm 15 well pre cast Novex 4-20% Tris-Glycine Gels (Invitrogen #XP04205BOX) and 30µl into 10 well pre cast gels. Ladder used was Color Protein Standard, Broad Range (11-245 kDa) (New England Biolabs, #P7712S). Lysates were run in 1x Tris-Glycine (Thermo) running buffer at 130V until the bands reached the bottom of the gel. Gels were transferred in transfer buffer at 4°C at 0.2 A onto a methanol activated polyvinylidene fluoride (PVDF) membrane (Immobilon-FL). Membranes were then stained with Ponceau (Sigma) dissolved in acetic acid to ascertain where the proteins were before cutting the membrane at required places. Membranes were then blocked in either 5% milk (Marvel) or 5% BSA if staining for phospho-proteins in Tris-buffered saline (TBS, prepared in house according to standard protocols) 0.1% Tween20 (TBST) solution for 1 hour at room temperature (RT)on a rolling shaker. Membranes were then washed once in TBST. Primary antibodies were diluted in either 5% milk TBST or 5% BSA TBST solution and incubated at 4**°**C overnight on a rolling shaker. 10% glycerol was added to primary antibody solutions to be able to reuse them post storing at -20**°**C. Membranes were washed 3 x 15 minutes in TBST. Membranes were then incubated for one hour at RT in either appropriate (mouse or rabbit) Horse radish peroxidase (HRP) - linked secondary antibodies (Cell Signaling) in TBST for chemiluminescent detection. Membranes were washed 3 x 5 minutes with TBST. Membranes were treated with ECL substrate (Thermo, #32106) and exposed using Azure Biosystems c600 western blot imaging system. Descriptions of antibodies used and dilutions can be found in Table 5. Western Blots were quantified using Image J and normalised to GAPDH unless otherwise stated.

**Table 5:** Primary antibodies used for Immunofluorescence (I.F) and Western Blot (W.B).

| Target | Antibody and species | Company and cat number | W.B dilution | I.F Dilution |
| --- | --- | --- | --- | --- |
| YAP/TAZ | Anti-YAP/TAZ (mouse) | Santa Cruz Biotechnologies, cat # A21235 | 1:1000 | 1:1000 |
| $\alpha$ -tubulin | Anti- α-tubulin (rat) | Bio-Rad, cat # MCA77G | | 1:1000 |
| GSK3β | Anti- | CST, cat # | | 1:300 |
| | GSK3β<br>(rabbit) | 12456T | | |
| pGSK3β | Anti-phospho<br>GSK3β<br>(Ser9)<br>(D85E12<br>(rabbit) | CST, cat #<br>5558T | | 1:400 |
| Golgi | Anti-GM180<br>(mouse) | Abcam, cat<br>#52649 |  | 1:500 |
| Beta-catenin | Anti- Beta-catenin<br>(mouse) | BD<br>Transduction<br>Laboratories,<br>cat # 610153 |  | 1:1000 |
| FOXO3A | Anti-FOXO3A<br>(75D8)<br>(rabbit) | CST, cat #<br>2497S |  | 1:200 |
| DOCK5 | Anti-DOCK5<br>(rabbit) | Frank et al.,<br>2017. | 1:500 |  |
| PHH3 | Anti- P-<br>Histone H3<br>(S10)<br>(mouse) | CST, cat #<br>9706S | 1:1000 |  |
| KIF5B | Anti-KIF5B<br>(rabbit) | Protein Tech,<br>cat #231632-1-AP | 1:1000 |  |
| PCNA | Anti- PCNA<br>(mouse) | Santa Cruz<br>Biotechnology,<br>cat #sc-56 | 1:1000 |  |
| GAPDH | Anti-GAPDH<br>(1D4)<br>(mouse) | Novus, cat #<br>NB300-221 | 1:2000 |  |
| Paxillin | Anti-Paxillin<br>(mouse) | BD transduction<br>labs # 610052 |  | 1:300 |

### Subcellular fractionation

Thermo’s Subcellular Protein Fractionation Kit for Cultured Cells (#78840) was used according to the manufacturer’s protocol to extract cytoplasmic, membrane, nuclear soluble, chromatin-bound and cytoskeletal proteins. Briefly, cells were harvested by trypsining them and centrifuging for 5 minutes. Cell pellets were washed with ice-cold PBS. Cells were counted so as to be able to use the corresponding ratios of Cytoplasmic Extraction Buffer (CEB), Membrane Extraction Buffer (MEB), Nuclear Extraction Buffer (NEB) and Pellet Extraction Buffer (PEB): CEB:MEB:NEB:PEB 200:200:100:100µl respectively. Fractions were stored at -80°C.

Protein concentrations of each subcellular fraction were measured using the Pierce BCA Protein Assay kit (Thermo). Samples were diluted in water (Sigma) to the lowest sample concentration to ensure the same amount of protein for each fraction was loaded into the gels. 4µl of NuPage LDS Sample Buffer (4x) were added to 16µl of each sample, which were then boiled for 10 minutes and spun down before freezing. Samples were then run as described in Western Blotting subsection.

### Mass spectrometry

#### Preparation and TMT labelling

LM2, MDA-MB-231, and hs578t cells were reverse transfected with *DOCK5* siRNA. LM2 and MDA-MB-231 were both plated at a density of 3.3 x 10^4^ cells/ml (low density, siRNA screening density) and 1.0 x 10^5^ cells /ml (high density) in T75 falcons due to the requirement of 3 million cells per sample. hs578t cells were plated at 5.0 x 10^4^ cells /ml. 48 hours later cells were collected by trypsinization and washed 3 x with cold PBS in clean mass spectrometry tubes. Samples were then flash frozen with 70% ethanol and dry ice. Cell pellets were dissolved in 150 μL lysis buffer containing 1% sodium deoxycholate (SDC), 100mM triethylammonium bicarbonate (TEAB), 10% isopropanol, 50mM NaCl and Halt protease and phosphatase inhibitor cocktail (100X) (Thermo, #78442) on ice, assisted with pulsed probe sonication for 15 sec. Samples were subsequently boiled at 90°C for 5 min on a thermomixer and sonicated for a 5 sec additionally. Protein concentration was measured with the Coomassie Plus Bradford Protein Assay (Pierce) according to manufacturer’s instructions. Aliquots containing 100μg of total protein were prepared for trypsin digestion. Samples were reduced with 5mM tris-2-carboxyethyl phosphine (TCEP) for 1h at 60 °C and alkylated with 10mM Iodoacetamide (IAA) for 30min in dark. Proteins were then digested by adding trypsin (Pierce) at final concentration 75ng/μ to each sample and incubating overnight. The resultant peptides were labelled with the TMT-10plex multiplex reagents (Thermo) according to manufacturer’s instructions and were combined in equal amounts to a single tube. The combined sample was then dried with a centrifugal vacuum concentrator.

#### High-pH Reversed-Phase Peptide Fractionation and LC-MS/MS Analysis

Offline high pH Reversed-Phase (RP) peptide fractionation was performed with the XBridge C18 column (2.1 x 150 mm, 3.5 μm, Waters) on a Dionex Ultimate 3000 High Performance Liquid Chromatography (HPLC) system. Mobile phase A was 0.1% ammonium hydroxide and mobile phase B was 100% acetonitrile, 0.1% ammonium hydroxide. The TMT labelled peptide mixture was reconstituted in 100 μL mobile phase A and was fractionated using a gradient elution method at 0.2 mL/min with the following steps: for 5 minutes isocratic at 5% B, for 35 min gradient to 35% B, gradient to 80% B in 5 min, isocratic for 5 minutes and re-equilibration to 5% B. Fractions were collected every 42 sec and vacuum dried.

LC-MS/MS analysis was performed on the Dionex Ultimate 3000 system coupled with the Q Exactive HF Orbitrap Mass Spectrometer (Thermo Scientific). Each peptide fraction was reconstituted in 40 μL 0.1% formic acid and 7 μL were loaded to the Acclaim PepMap 100, 100 μm × 2 cm C18, 5 μm 100 L trapping column at 10 μL/min flow rate. The sample was then subjected to a gradient elution on the EASY-Spray C18 capillary column (75 μm × 50 cm, 2 μm) at 45 °C. Mobile phase A was 0.1% formic acid and mobile phase B was 80% acetonitrile, 0.1% formic acid. The gradient separation method at flow rate 250 nL/min was as follows: for 90 min gradient from 5%-38% B, for 10 min up to 95% B, for 5 min isocratic at 95% B, re-equilibration to 5% B in 5 min, for 10 min isocratic at 5% B. The top 15 precursor ions between 350-1,850 m/z were selected with mass resolution of 120 k, AGC 3×10^6^ and max IT 50 ms for HCD fragmentation with isolation width 1.0 Th. Collision energy was set at 35% with AGC 1×10^5^ and max IT 100 ms at 60k resolution. Targeted precursors were dynamically excluded for 30 seconds.

#### Database search and protein quantification

The SequestHT search engine was used to analyse the acquired mass spectra in Proteome Discoverer 2.2 (Thermo Scientific) for protein identification and quantification. The precursor mass tolerance was set at 20 ppm and the fragment ion mass tolerance was set at 0.02 Da. Spectra were searched for fully tryptic peptides with a maximum of 2 miss-cleavages. TMT6plex at N-terminus/K and Carbamidomethyl at C were defined as static modifications. Dynamic modifications included oxidation of M and Deamidation of N/Q. Peptide confidence was estimated with the Percolator node. Peptide False Discovery Rate (FDR) was set at 0.01 and validation was based on q-value and decoy database search. All spectra were searched against reviewed UniProt human protein entries. The reporter ion quantifier node included a TMT 10plex quantification method with an integration window tolerance of 15 ppm and integration method based on the most confident centroid peak at the MS2 level. Only unique peptides were used for quantification, considering protein groups for peptide uniqueness. Peptides with average reporter signal-to-noise >3 were used for protein quantification.

Abundance values were scaled within each cell line and the log2 ratio of abundance in si*DOCK5* containing samples to mock transfected samples was calculated. Log2 ratios above 0.5 and below -0.5 and permutation FDR <0.05 (t-test) were considered to be proteins which were significantly up (∼40% minimum increase) and down regulated (∼40% minimum decrease) respectively. Statistical analysis of proteomics data was done in the Perseus software(Tyanova et al., 2016).

### Microscopy

#### Immunofluorescence and confocal microscopy

All performed as described for siRNA screening. Antibodies and their dilutions can be found in Tables 5 and 6. Unless otherwise stated cells were imaged using an automated Opera Quadruple Enhanced High Sensitivity (QEHS) spinning-disk confocal microscope (PerkinElmer) with 20x air lens. Lasers and their corresponding filters used were: 405(450/50), 561 (600/40), 488 (540/751), 640 (690/50).

**Table 6:** Secondary antibodies used for Immunofluorescence (I.F) and Western Blot (W.B).

| Secondary antibody/dyes | Company | Dilution |
| --- | --- | --- |
| Phalloidin | Invitrogen, cat # A12379 | 1:1000 |
| Hoechst | Sigma Aldrich, cat # 33258 | 1:1000 |
| AlexaFluor 647 goat anti-mouse | Life Technologies, cat # A21235 | 1:1000 |
| AlexaFluor 568 goat anti-rat | Life Technologies, cat # A11077 | 1:1000 |
| AlexaFluor 647 goat anti-rabbit | Life Technologies, cat # A21245 | 1:1000 |
| AlexaFluor 568 goat anti-rabbit | Life Technologies, cat # A11036 | 1:1000 |
| Anti-mouse IgG, HRP-linked antibody | CST, cat #7076S | 1:2000 |
| Anti-rabbit IgG, HRP-linked antibody | CST, cat #7074S | 1:2000 |

#### Focal adhesion dynamics and Total Internal Reflection Fluorescence Imaging (TIRF)

LM2 CAAX-GFP cells were transfected with either just Talin mApple plasmid or both Talin mApple and si*DOCK5,* si*RAC,* si*RHOA*, or si*CDC42* as stated in the transfections section. 35mm glass Mat-tek dishes were coated with 10µg/ml Fibronectin (human plasma, Sigma #F0895) for 1 hour. Fibronectin was then aspirated and Mat-tek dishes were washed 3 x PBS and left to dry. 48 hours after transfection, LM2 CAAX-GFP cells were plated at 1.5 x 10^5^ cells /ml in 1ml of Liebnevitz medium (10% FBS, 1% penicillin/streptomyocin). Cells were live imaged using the TIRF (3i) approximately 1 hour after seeding. Lasers used were 647 And 488. Images were taken either every 3 minutes or every 1.5 minutes. Images were acquired using Slidebook software (3i). Movies were processed to be the same length and analysed using the ‘Focal Adhesion Analysis Server (FAAS)’ (Berginski and Gomez, 2013). Movies containing the same number of time frames were compared.

We displayed the data using estimation statistics using a website generated by Ho et al., 2019, as we think that by showing the effect size we can visualise and understand this type of data better.

#### Spreading assay and Super-Resolution via Optical-Reassignment (SoRa) microscopy

Cells pertaining to experiments imaged using the SoRa were fixed and stained with a slight variation to those from RNAi screens. Briefly, cells were reverse transfected as described in the transfection section in 6 well plates, and plated on 35mm glass Mat-tek dishes coated with 10µg/ml Fibronectin as described for TIRF, at a density of 1.5 x 10^5^ cells /ml 48 hours after transfection. They were fixed 3 hours after plating with paraformaldehyde in PBS (4% final concentration), permeabilised with 0.1% Triton-X-100 in PBS for 10 minutes, and blocked with 0.5% BSA dissolved in PBS with 0.3ml of glycine for 1 hour. Antibody incubations were performed in 0.5% BSA. Primary antibodies were added at RT for 1.5 hours. Secondary antibodies were added for 1.5 hours at RT. Hoechst was used as a DNA stain and left for 15 minutes at the end at RT. Cells were washed 3 x with PBS between each step and were left in 200µl of PBS. Where antibodies used were close in species such as mouse and rat, they were added sequentially to avoid any possible bleedthrough. Antibodies and their dilutions can be found in Tables 5 and 6. When drugs were used, they were added in 500µl on top of the 2ml solution of cells immediately after plating, with final concentrations as shown in Table 2. Cells were imaged using the SoRa (3i) microscope.

### Wound heal assay

Cells were reverse transfected as previously described and were plated in a 96 well Perkin Elmer Cell Carrier Ultra plate at a density of 2.7 x 10^6^ cells/ml. 48 hours later a 10μM pipette tip was used to scratch the monolayers from top to bottom of the well, resulting in a vertical cell-free gap or wound. Medium was removed and wells were carefully washed with warm PBS so as to remove debris. 100 μL of fresh medium was them added to all wells. Imaging began as soon as the wound was created, using a Basic Confocal Spinning Disk Microscope (3i). Images were taken every 10 minutes for 16 hours. Images were processed using ImageJ. We quantified Golgi orientation in cells that had their Golgi within 180 degrees of the wound and quantified the angle of orientation by drawing a line from the position of the Golgi towards the wound and another parallel to the wound (Fig. 7e, ii) as in Zhang et al., 2008(Zhang et al., 2008). A 90 degree angle is indicative of Golgi facing toward the wound and indicative of front-rear polarity.

### Growth curve assay

Cells were reverse transfected in 6-well plates and plated at a density of 3.3 x 10^4^ cells/ml per well, with three replicates per condition. When drugs were used one plate containing mock transfected cells and *DOCK5* siRNA would be treated with DMSO, and another with the drug. Binimetinib (MEKi) and FAKi (PF-573288) were added to a final concentration of 10µM and 2 μM respectively. Drugs and DMSO were added in 500 μl DMEM 48 hours after transfection. Plates were incubated in the IncuCyte S3 Live-Cell Analysis System (Sartorius) at 37**°**C and 5% CO_2_ and imaged at 6 hour time points for a minimum of 156 hours (6.5 days). A minimum of 9 fields of view were imaged per well. Analysis of confluence over time was carried out using IncuCyte software (Sartorius). To compare confluence between conditions at a point where mock transfected cells reached 50% or 100% confluency, each well was normalized to the mock transfected wells for each plate, prior to statistical analysis. Representative growth curves are shown per plate with the average of three wells per condition.

### Statistical analyses

Statistical analyses performed for each assay are indicated in text and Figure legends. Most analyses were carried out using Prism 8 (GraphPad). PCA analysis was performed in Jupyter Notebooks (Kluyver et al., 2016). Hierarchical clustering and heat maps were made with Morpheus Broad Institute Software (https://software.broadinstitute.org/morpheus).

## Acknowledgements

We thank Nicola Ferrari (Astex Pharmaceuticals, UK) for the CAAX-GFP construct, Aaron Farrugia for 2-photon imaging (ICR, London, UK), and Vicky Bousgouni (Dynamical Cell Systems Team, ICR, London, UK) for assistance with the RNAi library array and helpful comments on the manuscript. We thank Oliver Rocks (Max-Delbrück-Center for Molecular Medicine, Berlin, Germany) for the DOCK5-YFP construct and Steen Hansen (Boston Children’s Hospital, Boston, USA) for the DOCK5 antibody. We thank Frankie Butera for helpful comments on the manuscript. This work was funded by a Cancer Research UK and Stand Up to Cancer UK Programme Foundation Award to C.B. (C37275/1A20146). The work of T.I.R and J.S.C was funded by the CRUK Centre grant with reference number C309/A25144.

## Author Contributions

P.P.-V. contributed to experimental design, carried out the experiments, performed image and data analysis, wrote and prepared the manuscript. M.A.-G. optimized RNAi screening methodology, performed experiments, data and image analysis. T.I.R, J.S.C performed mass spectrometry and proteomic analysis. C.B contributed to experimental design, imaging, and manuscript writing and preparation.

## Figure legends

**Supplementary Fig. 1. Validation of *DOCK5* siRNA knockdown.** (a) Graph depicting log10 ratio of nuclear to ring region YAP/TAZ versus cell number for MDA-MB-231 (light blue), LM2 (dark blue), SUM159 (orange), hs578t (yellow), T47D (pink), and MCF10A (orange). Representative images of hs578t, SUM159, MCF10A, T47D, MDA-MB-231, and LM2 at low (3.3 x 10^4^ cells/ml) and high (1.3 x 10^5^ cells/ml) densities, stained for YAP/TAZ. As well as 2.7 x 10^5^ cells/ml for MCF10A and T47D. n = 4 wells/condition. Scale bar is 50µm. (b) Z-scores per well for YAP/TAZ ratio for siGENOME siRNA screen in LM2. Above 1.5 (red dotted line) are siRNAs which are hits for high YAP/TAZ ratio (YAP/TAZ inhibitors). Below -1.5 (red dotted line) are siRNAs which are hits for low YAP/TAZ ratio (YAP/TAZ activators). Four individual replicates for si*DOCK5* as hits for low YAP/TAZ ratio are shown. (c) Graphs showing YAP/TAZ ratio and YAP/TAZ levels upon transfection with DOCK5 deconvoluted and pooled siRNAs for OTP and siGENOME. Mean ± sd (n > 3 wells, 1000 cells/well. ANOVA, *P < 0.1** P < 0.01, *** P < 0.001**** P < 0.0001). (d) qRTPCR showing *DOCK5* mRNA levels in mock versus si*DOCK5* transfected cells. Mean ± s.d (n = 3 biological repeats, at least 3 technical replicates each, Student’s t-test). (e) Validation of DOCK5 siRNA knockdown via qRTPCR. *DOCK5* mRNA levels following *DOCK5* knockdown with deconvoluted siRNAs for DOCK5 OTP5 to 8 and in pooled combination (OTP pool), as well as siG1-4 and in pooled combination (siGENOME pool). Mean ± sd (n = 4 wells, ANOVA, **** P < 0.0001). (f) (g) Representative western blots of subcellular fractions for mock and si*DOCK5*. 1: nuclear fraction; 2: membrane fraction; 3: cytoplasm fraction; 4: chromatin bound fraction, 5: cytoskeletal fraction. M is mock, D5: si*DOCK5*. (f) Blotted for YAP and TAZ. Loaded 5ng of protein for all except for cytoskeleton (2.5ng). (g) Blotted for DOCK5. WC L is indicative of whole cell lysate. Below, quantification of DOCK5 protein levels upon *DOCK5* knockdown for whole cell lysates. Mean ± SEM (n = 3, Student’s t-test with Welch’s correction, ** P < 0.01). (h) Same subcellular fractions as in (e) blotted for PH3B, KIF5B, GAPDH, and PCNA as controls for each fraction: chromatin bound, cytoskeleton bound, whole cell, nuclear respectively. Pie charts depicting the distribution of (i) PCNA, (ii) KIF5B, (iii) PH3B, in each fraction.

**Supplementary Fig. 2. Z-scores for YAP/TAZ ratio in siGENOME and OTP screens in MDA-MB-231.** Z-scores per well for YAP/TAZ ratio for siGENOME siRNA screen in MDA-MB-231. Above 1.5 (red dotted line) are siRNAs which are hits for high YAP/TAZ ratio (YAP/TAZ inhibitors). Below -1.5 (red dotted line) are siRNAs which are hits for low YAP/TAZ ratio (YAP/TAZ activators). Four individual replicates for si*DOCK5* as hits for low YAP/TAZ ratio are shown.

**Supplementary Fig. 3. RHOA, RAC1, YAP, TAZ depletion do not affect growth rates.** Growth curves for mock and (a) si*DOCK5*, (b) si*CDC42* (c) si*RHOA*, (d) si*RAC1*, (e) si*YAP* (f) si*TAZ* and (g) si*YAP* + si*TAZ* transfected cells. Mock and siRNA transfected wells represented as circles and squares respectively. Experiments performed in triplicate, representative experiment shown with n = 3 wells per condition. Averages per well are shown.

**Supplementary Fig. 4. Shape filtering in LM2.** (a) (b) PCA of single cell dataset comprised of texture and morphological features of populations which were either mock transfected (a) or transfected with siRNAs which scored the highest for enriching for a particular shape (b). Each data point represents a cell and is colour coded by the shape into which it has been classified. Spindly in red, round large in green, triangular in blue, fan in black, and small round in purple. Black box represents 1 standard deviation of the mean for PC1 and PC2 of (b), projected on (a). Single cells within black box shown in (a) were defined as ‘normal’ in the new classifiers. (c) PCA of single cell dataset comprised of texture and morphological features of siRNA transfected population, after running new classifiers on same data, removing cells classified as ‘normal’. Ellipses represent 95% confidence intervals for each shape.

**Supplementary Fig. 5. Beta catenin upregulation in response to LiCl treatment and increased nuclear FOXO3A upon Uprosertib treatment.** (a) Representative images of mock transfected cells treated with DMSO, 5mM LiCl, or 2μM final Uprosertib. Stained for FOXO3A (yellow), YAP/TAZ (red), and merge including F-actin (green) and DNA (blue). Scale bar is 50 µm. Arrows point to nuclear FOXO3A and YAP/TAZ which are higher and lower respectively in Uprosertib. (b) Quantification of FOXO3A nuclear to cytoplasmic ratio (log) shown in a. Mean ± sd are shown (n > 3 wells, 1000 cells per well. ANOVA, **** P < 0.0001). (c) Representative images of mock transfected cells treated with DMSO, 5mM LiCl, 2μM final Uprosertib. Stained for GSK3β (yellow), βcatenin (red), and merge including F-actin (green) and DNA (blue). Scale bar is 50 µm. Arrows indicate GSK3β at leading edge and increased beta catenin. (d) Quantification of beta catenin shown in b. Mean ± sd are shown (n > 3 wells, 1000 cells per well. ANOVA, **** P < 0.0001).

**Supplementary Fig. 6. Automated segmentation for Collagen Invasion Quantification.** Images for mock and si*DOCK5* transfected cells shown at planes quantified: bottom plane 0μm and middle plane 30μm. As well as intermediate planes 15μm, and 45μm. Cells selected indicate cells that the software has identified to be in that particular plane (green) following a filtering process. Those that are not considered to be in that plane are in red. Shows that cells which are counted in middle plane are not spanning other planes. Scale bar is 50 µm.

**Supplementary Movie 1.** Movie showing LM2-CAAX GFP cell overexpressing Talin mApple (white). Total Movie time is 63 frames with each frame taken every 3 minutes. Exported at 7 frames per second.

**Supplementary Movie 2.** Movie showing LM2-CAAX GFP cell transfected with si*DOCK5* and overexpressing Talin mApple (white). Total Movie time is 63 frames with each frame taken every 3 minutes. Exported at 7 frames per second.

**Supplementary Movie 3.** Movie showing LM2-CAAX GFP cell transfected with si*CDC42* and overexpressing Talin mApple (white). Total Movie time is 50 frames with each frame taken every 1.5 minutes. Exported at 7 frames per second.

**Supplementary Movie 4.** Movie showing LM2-CAAX GFP cell transfected with si*RAC1* and overexpressing Talin mApple (white). Total Movie time is 50 frames with each frame taken every 1.5 minutes. Exported at 7 frames per second.

**Supplementary Movie 5.** Movie showing LM2-CAAX GFP cell transfected with si*RHOA* and overexpressing Talin mApple (white). Total Movie time is 50 frames with each frame taken every 1.5 minutes. Exported at 7 frames per second.

**Supplementary Movie 6.** Movie showing LM2-CAAX GFP cell overexpressing Talin mApple (white) and treated with Uprosertib. Total Movie time is 50 frames with each frame taken every 1.5 minutes. Exported at 7 frames per second.

