## Supplementary figures for "Multiplexed quantitative screens of single cell shape and YAP/TAZ localisation identify DOCK5 as a coincident detector of polarity and adhesion during migration"

Supplementary Figure 1 Validation of DOCK5 siRNA knockdown.

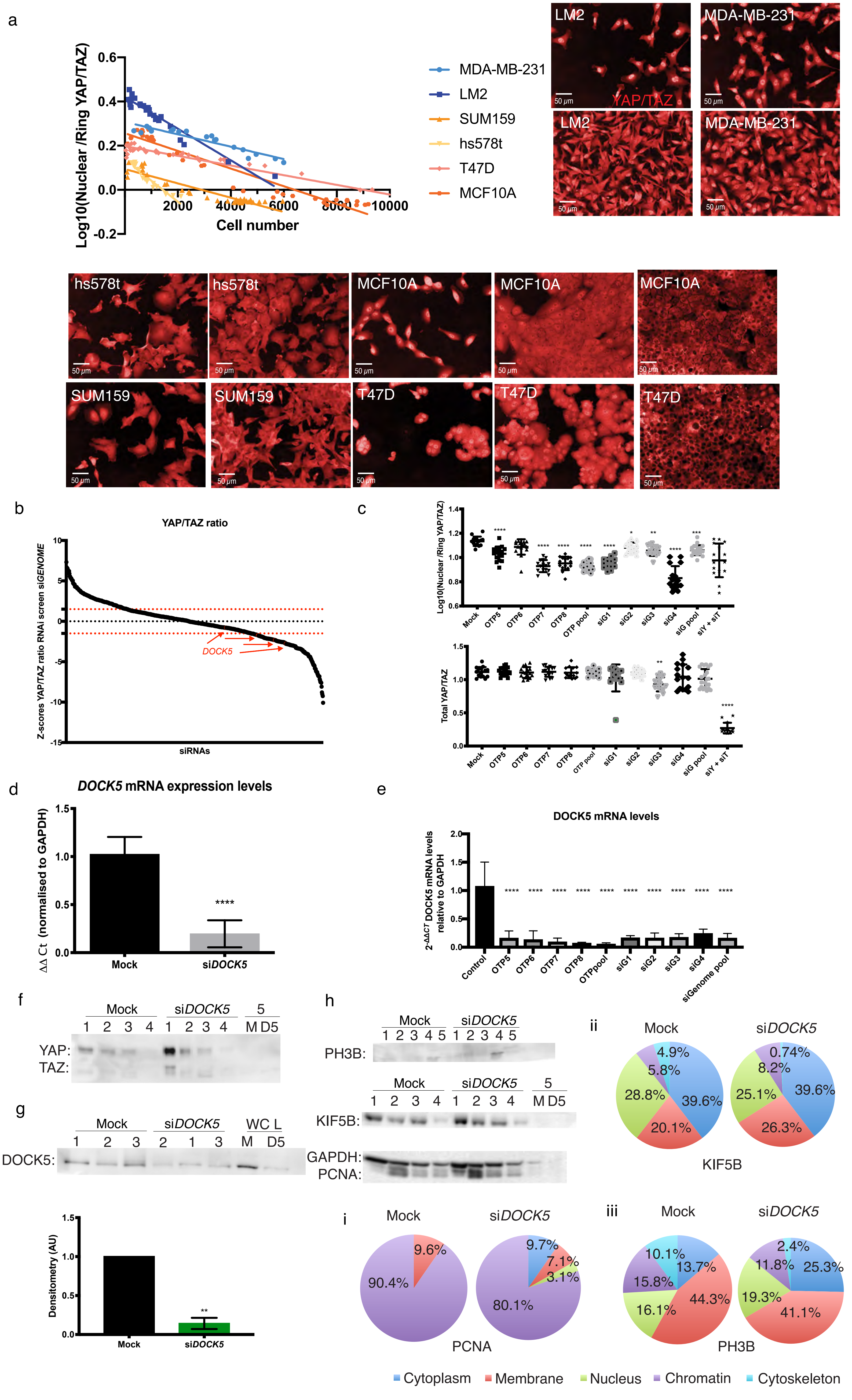

Supplementary Fig. 2. Z-scores for YAP/TAZ ratio in siGENOME and OTP screens in MDA-MB-231.

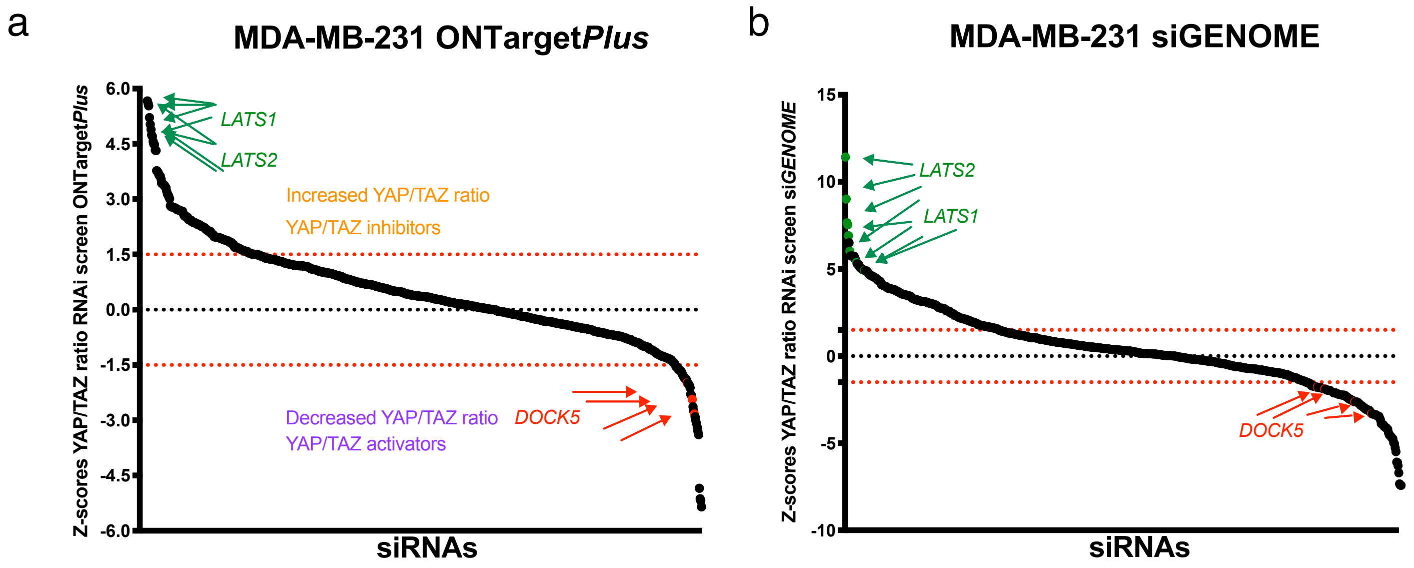

Supplementary Figure 3 RHOA, RAC1, YAP, TAZ, depletion do not affect growth rates

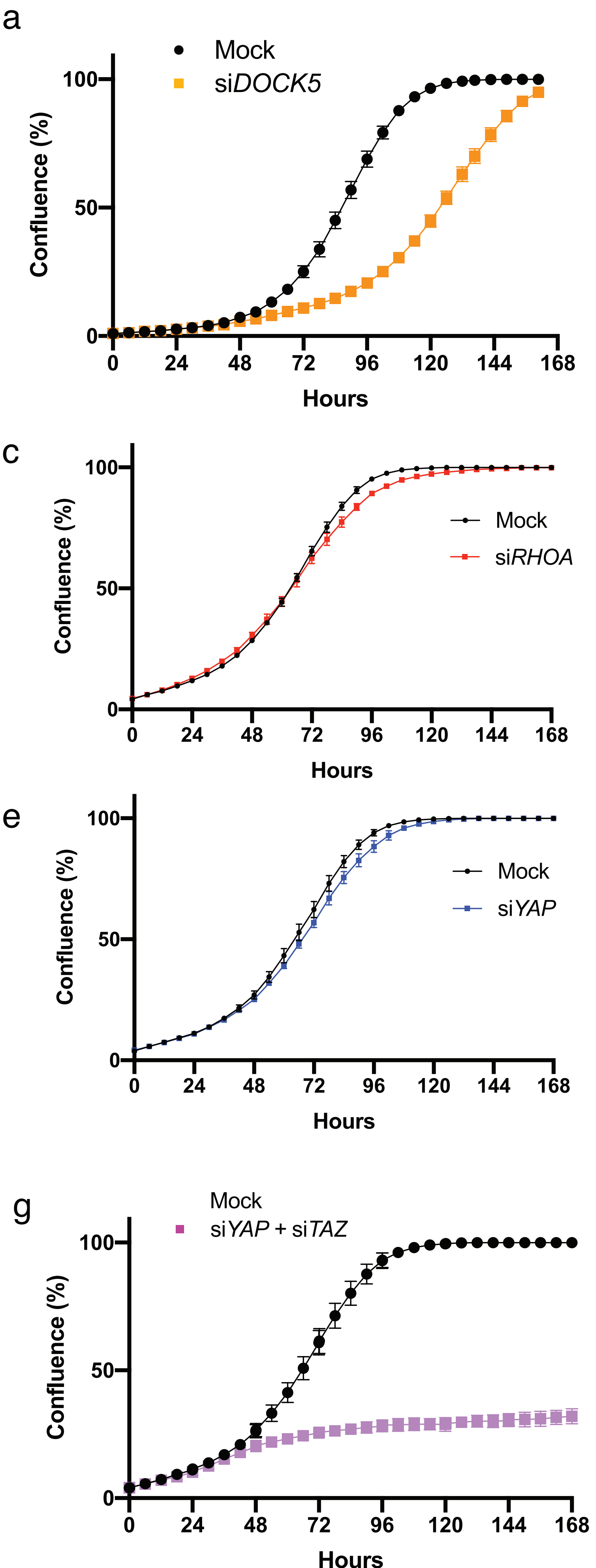

### Supplementary Figure 4. Shape filtering in LM2

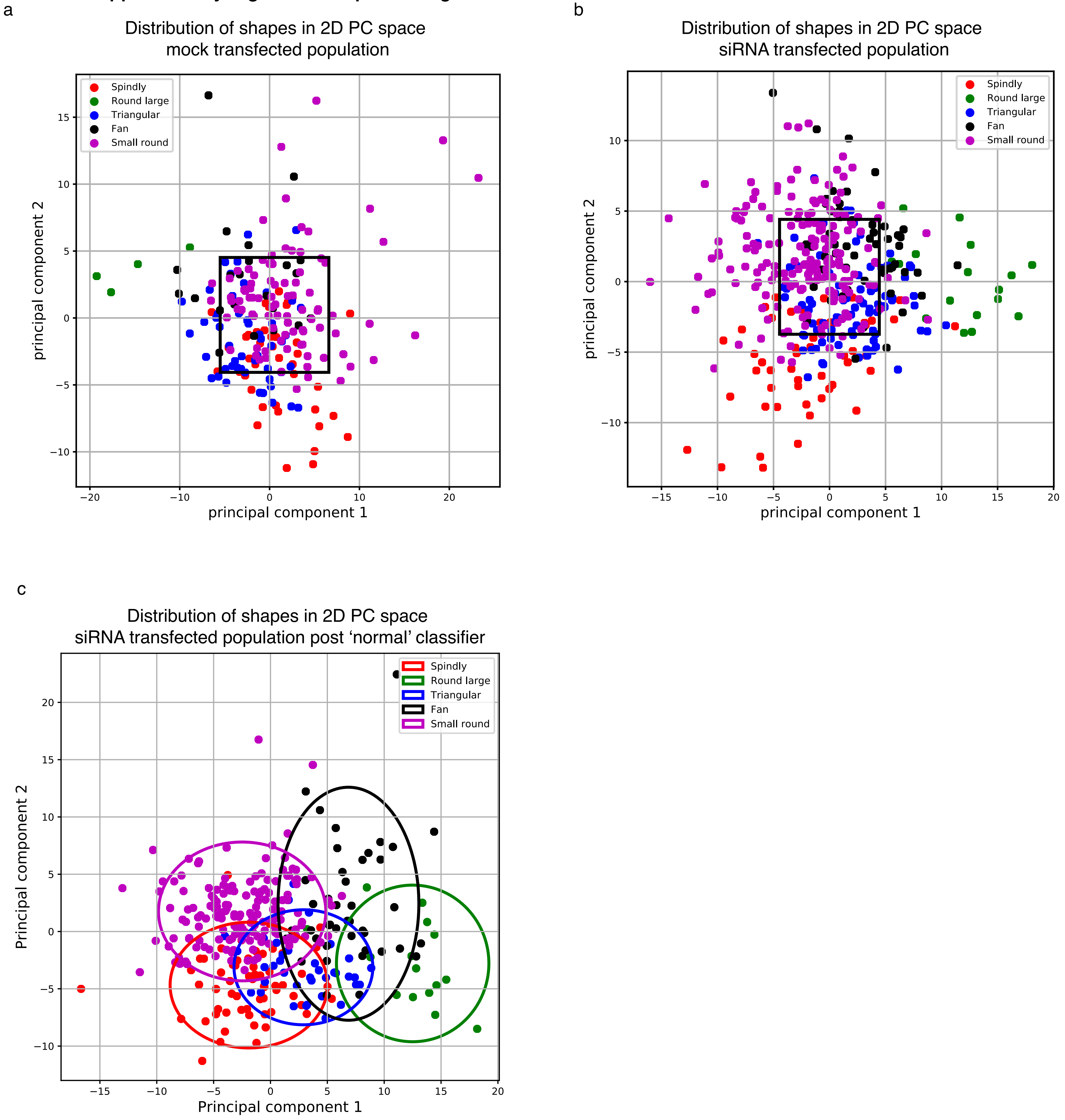

Supplementary Figure 5 Beta catenin upregulation in response to LiCl treatment and increased nuclear FOXO3A upon Uprosertib treatment

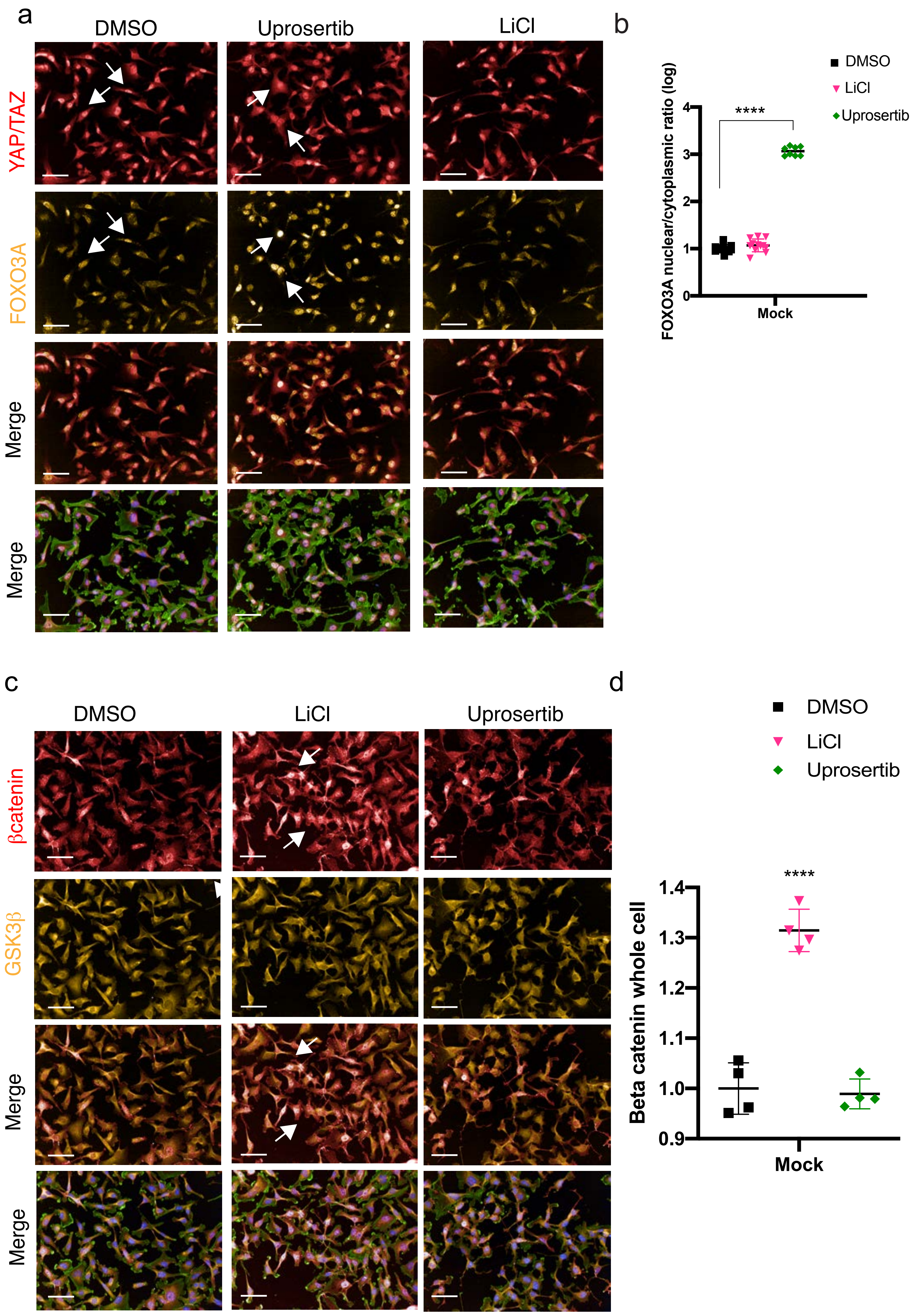

Supplementary Figure 6. Automated segmentation for Collagen Invasion Quantification

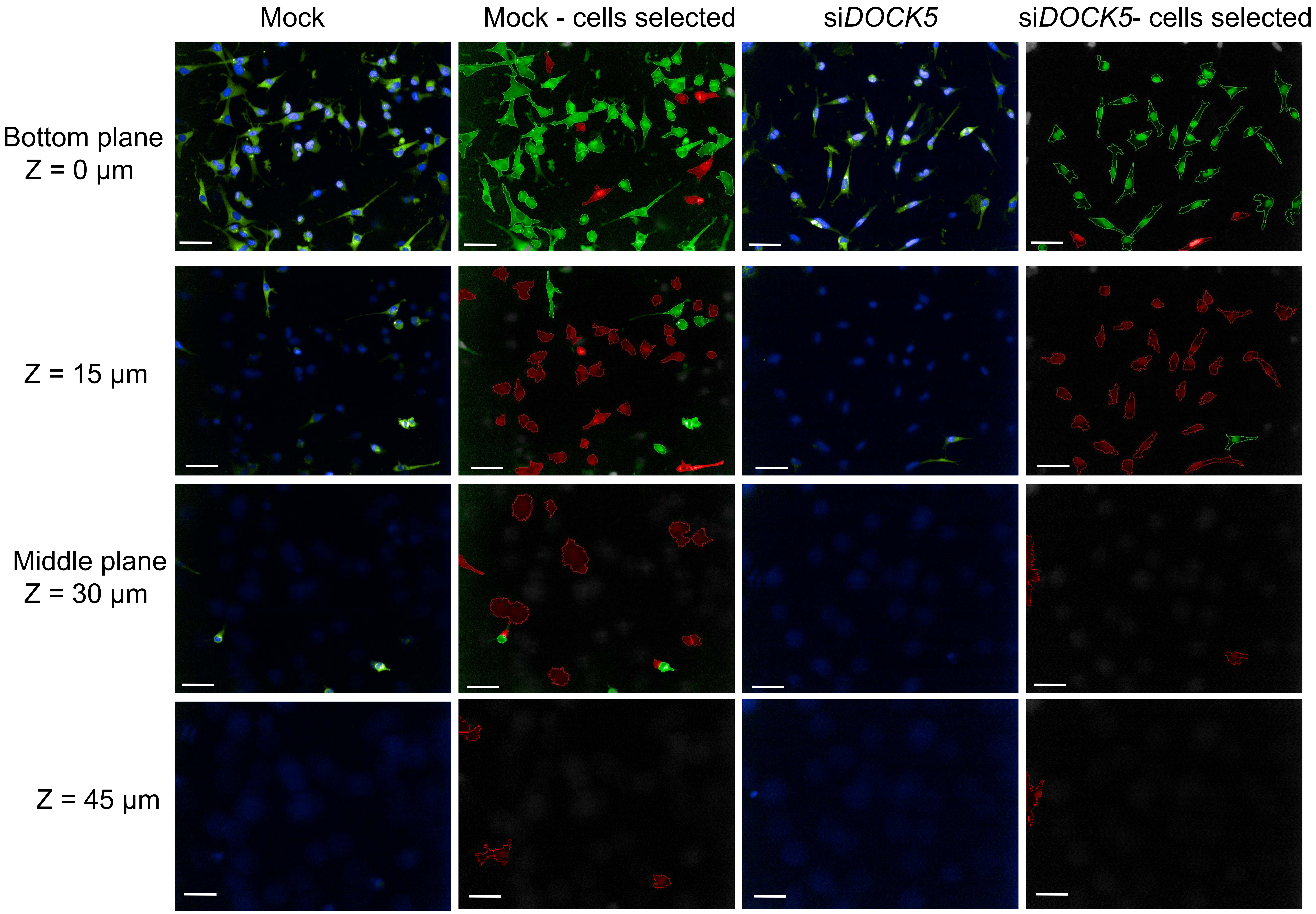
